# Ribosomal RNA tentacles are targets of free radical damage in mammalian cells during oxidative and inflammatory stress

**DOI:** 10.1101/2025.09.11.675630

**Authors:** Aaron M. Fleming, Justin C. Dingman, Cynthia J. Burrows

## Abstract

Chemical damage to ribosomal RNA (rRNA) during oxidative or inflammatory stress can impact protein synthesis. Human cells were exposed to a H_2_O_2_ titration series to induce oxidative stress or to tumor necrosis factor-α to induce inflammation over a time course followed by RNA direct nanopore sequencing of cytosolic and mitochondrial rRNAs. The guanosine (G) oxidation sites and deamination of adenosine to inosine (A-to-I) and cytidine to uridine (C-to-U) lesion sites were revealed by changes in the base-called data. Both stressors induced G oxidation in cytosolic rRNA, whereas mitochondrial rRNA was less oxidatively modified. Nitrosative stress generated during inflammation resulted in deamination lesions in rRNAs in both compartments. Inspection of highly modified sites showed the GC-rich tentacles in the 28S rRNA sequence were hotspots for G oxidation and C deamination in the cytosolic ribosome. Outside of tentacles, lesions were generally found on nucleotides on the ribosome surface exposed to solvent, where diffusible reactive species exist. The minimalist structure of the mitochondrial ribosome compared to the cytosolic ribosome alters the reaction patterns observed to target nucleotides on the surface or in functionally relevant regions. These patterns support the hypothesis that tentacles in cytosolic ribosomes direct reactive oxygen and nitrogen species away from the catalytic core to maintain ribosome activity during stress, while the mitochondrial ribosome is damaged in regions that can deactivate protein synthesis. The results provide molecular insight into metabolic dysfunction during oxidative and inflammatory stress and suggest a new function for the GC-rich tentacles that have evolved in mammalian cells.

**Significance Statement:** Infection and injury trigger a cellular inflammatory response resulting in the release of free radical species capable of DNA and RNA damage. We used RNA direct nanopore sequencing to map oxidized guanosine sites in human ribosomal RNA via base-calling error analysis. Reactive nitrogen species derived from peroxynitrite result in deamination reactions, principally cytidine to uridine and adenosine to inosine, which can be directly read by nanopore sequencing. Importantly, cytosolic ribosomes behave very differently than mitochondrial ribosomes; the latter rRNA is somewhat protected from G oxidation by high levels of bicarbonate as well as a protein-coated ribosome structure. In contrast, cytosolic rRNA tentacles are hotspots for both G oxidation and C and A deamination, which might explain their evolved function.

## Introduction

Nucleic acids are damaged by reactive oxygen and nitrogen species (RONS) generated in cells experiencing oxidative stress or inflammation. Historically, studies of DNA damage have far outpaced investigations into RNA damage, largely because DNA lesions can cause mutations implicated in disease and aging (1). However, RNA damage by RONS is more prevalent than DNA damage (2, 3). This increased susceptibility arises from several factors, including the fact that cells contain ∼5-fold more RNA than DNA (∼6 pg DNA vs. ∼30 pg RNA per human cell) (4); RNA is distributed across nearly all cellular compartments, unlike DNA; RNA secondary and tertiary structure is more solvent-exposed than DNA; and repair pathways for RNA damage have not been described, while similar pathways are highly evolved for DNA damage repair (1). Collectively, these features render RNA more prone to diffusible RONS-mediated free radical damage, and the damage may persist in cellular RNAs with low turnover rates. Within the human transcriptome, ribosomal RNA comprises >80% of the total mass and has a low recycling rate in cells (*t*_1/2_ > 3 days) (4). Consequently, rRNA is expected to bear the brunt of RONS-induced injury that may persist in cells, providing a trackable molecular signature of the RONS species generated during cellular stress.

The research community’s understanding of RONS-mediated RNA lesions has predominantly focused on oxidatively derived damage. The primary endogenous reactive oxygen species generated in cells results from incomplete metabolism, generating superoxide (O_2_^•-^) as an unwanted byproduct. Cells detoxify O_2_^•-^enzymatically via a dismutation reaction, furnishing H_2_O_2_. The fate of H_2_O_2_ is enzymatic clearance, yielding O_2_ and H_2_O via a disproportionation reaction, or it can undergo deleterious reactions that damage the cell, principally the Fenton reaction, in which H_2_O_2_ reacts with the redox-active or “labile” Fe^2+^ pool. Until recently, it was proposed the cellular Fenton reaction yields hydroxyl radical (HO^•^) or a ferryl species (Fe=O^2+^), depending on the pH and Fe^2+^ ligand (5); however, this view shifted with the recognition that bicarbonate (HCO_3_^-^), present in cells at >20 mM, is a Fe^2+^ ligand that influences the outcome of the reaction. The Fe^2+^-bicarbonate complex catalyzes the Fenton reaction with H_2_O_2_ to yield carbonate radical anion (CO_3_^•-^) rather than HO^•^/Fe=O^2+^ (6), and this finding was substantiated in human cells (7). This subtle but profound shift in the identity of the reactive species is critical because HO^•^ and Fe=O^2+^ are highly indiscriminate oxidants reacting with all nucleobases and the sugar-phosphate backbone of DNA or RNA (8, 9). In contrast, CO_3_^•-^ selectively oxidizes the guanine heterocycle in DNA to produce a defined set of products (10, 11).

The products of nucleic acid oxidation by CO_3_^•-^ were best determined in DNA, and this knowledge should translate to RNA, with the caveat that the relative product yields may differ due to structural and base-pairing effects influencing the product distribution (12). The major two-electron oxidation product of guanosine (G) is 8-oxo-7,8-dihydroguanosine (rOG; Figure 1A) (13); However, OG is more labile toward a second oxidation to yield the ribonucleosides of spiroiminodihydantoin (Sp) or 5-guanidinohydantoin (Gh), in which their relative yields are determined by the reaction context and pH (Figure 1A) (12, 14). Alternatively, CO_3_^•-^ can oxidize the DNA base guanine at C5 to yield a 2-iminohydantoin 2′-deoxynucleoside that should also occur in RNA (15). Another potential product is oxidation-initiated hydrolysis of G to yield 2,6-diamino-4-hydroxy-5-formamidopyrimidine ribonucleoside (Fapy-G) (Figure 1A) (16, 17). Routinely, OG is found in the transcriptome of cells experiencing stress at higher levels than its 2’-deoxy analog in the genome of the same cells (2, 3, 18). Moreover, OG is found at high levels in rRNA in unstressed prokaryotic cells (∼1 per 10^3^ rGs), which rises with H_2_O_2_ addition to the culture; and lastly, Gh and Fapy-G were found in prokaryotic total RNA under extreme oxidative stress conditions (17). The lesions Sp and 2Ih remain elusive in cellular RNA samples. Sequencing studies for OG-containing strands in mammalian mRNA and microRNA have shown preferential enrichment of the damaged nucleotide in specific transcripts within these RNA pools (19, 20), suggesting G oxidation exhibits structural or sequence-based selectivity in cells. Cellular responses to oxidized mRNA include increased ribosome stalling, activation of no-go decay pathways, and misalignment of tRNA anticodons with mRNA codons, which can alter the resulting protein sequence (21, 22). Thus far, we know G oxidation is prevalent in the transcriptome, and the lesions impact biological processes, although deficiencies in our knowledge remain, such as, where do the lesions occur?

**Figure 1.**
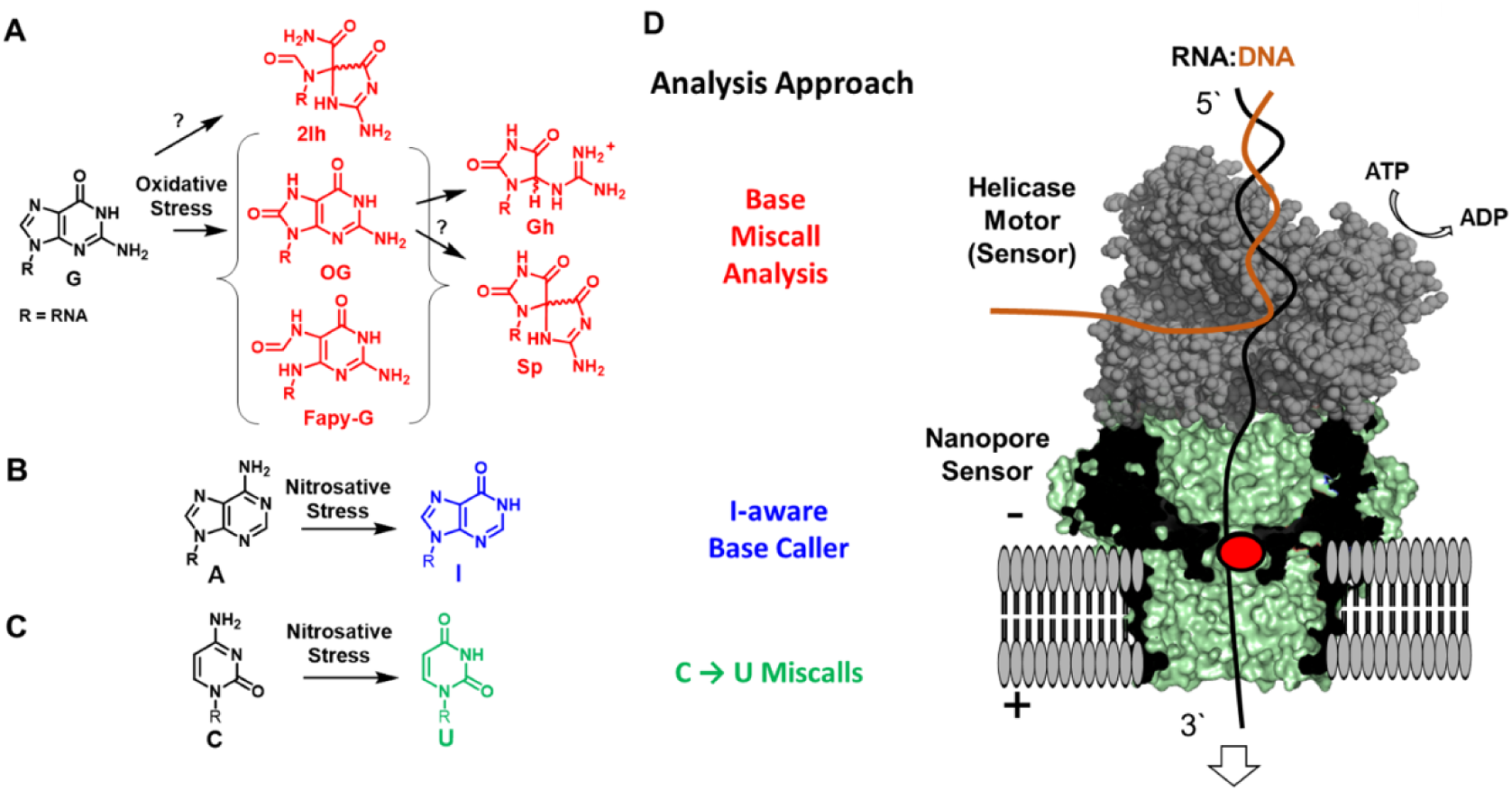
Oxidative and inflammatory stress generates RONS that can modify RNA that is detectable by RNA direct nanopore sequencing. Scheme of RNA modification products for (A) G oxidation, (B) A deamination, and (C) C deamination. (D) Schematic of RNA nanopore sequencer and the data analysis approaches to sequence the RNA lesions.

Inflammation produces a reactive species landscape that differs from purely oxidative stress leading to different patterns of DNA damage; however, the chemistry of these species to damage RNA remains a topic that has received less attention. Inflammatory processes induce the enzymatic production of O_2_^•-^and nitric oxide (^•^NO), which either rapidly combine to form peroxynitrite (ONOO^-^) (23), or the O_2_^•-^is enzymatically converted to H_2_O_2_ that reacts to yield CO_3_^•-^ via the bicarbonate-Fenton reaction (6, 7). In the presence of dissolved CO_2_, ONOO^-^decomposes to yield both CO_3_^•-^ and nitrogen dioxide radical (^•^NO_2_) (13, 24). As noted above, CO_3_^•-^ selectively oxidizes G (Figure 1A). Nitrogen dioxide radical is proposed to react further to form the potent nitrosating agent dinitrogen trioxide (N_2_O_3_) (25). In DNA, N_2_O_3_ induces deamination of 2′-deoxycytidine to 2′-deoxyuridine (dC-to-dU) in a twofold greater yield than deamination of 2′-deoxyadenosine to 2′-deoxyinosine (dA-to-dI) (26). Monitoring RNA deamination presents a challenge, particularly for LC-MS/MS of nucleosides, because uridine is a canonical nucleoside in RNA, and I is also enzymatically installed by ADAR enzymes (27), resulting in a need to decouple I as a form of RNA damage from I written in RNA with intent. In a test tube reaction, I is found from nitrosative stress imposed on model RNAs (28). In a mouse model experiencing ^•^NO overproduction, I in total RNA was only found to increase in tissues of the spleen, while the mechanism of introduction, being either enzymatic or RONS associated, could not be definitively decoupled (29). In summary, our understanding of nitrosative stress inducing lesions in cellular RNA remains limited.

In the present study, we exposed human cells to defined oxidative or inflammatory stress conditions and performed RNA direct sequencing of cytosolic and mitochondrial rRNAs using a commercial nanopore sequencer (Figure 1D). The G oxidation sites were detected through changes in miscall frequencies at G reference positions before and after stress. The C-to-U deamination was monitored by increases in U miscalls at reference C positions following stress exposure. The A-to-I deamination sites were identified through changes in I calls using the I-aware base call model in the Dorado nanopore signal base caller. The RNA direct nanopore sequencing revealed that the GC-rich tentacles of human cytosolic 28S rRNA are hotspots for oxidative and nitrosative damage, while the pattern of mitochondrial ribosome damage occurs most commonly on surface-exposed sites in the rRNAs. The biological consequences of damage to these regions of the ribosome are discussed.

## Results

### Inflammation-induced RONS oxidatively modify RNA at G to yield OG

The HEK293T cell line was chosen due to the availability of established protocols to guide experimental design (30). Cells were cultured in DMEM under 5% CO_2_ and 80% humidity, maintaining HCO_3_^-^levels at ∼20 mM. At ∼50% confluence, cells were treated with 25 ng/mL tumor necrosis factor-α (TNF-α) for 1–72 h; for treatments exceeding 24 h, medium and TNF-α were refreshed every 24 h. The cytokine TNF-α was selected to induce inflammation, as it is a common mediator of this cellular program over both short and long durations (30). Cell viability was monitored via a trypan blue exclusion assay, with >70% remaining viable at 72 h (Figure 2A). Oxidative stress was assessed using the fluorescence activation of 2,7-dichlorodihydrofluorescein diacetate (DCFH-DA) upon oxidation. The DCFH-DA assay showed maximal fluorescence between 2.5–12 h, which declined from 24–72 h, with final stabilization at ∼2-fold above untreated cells (0 h; Figure 2B). These results confirm cell viability during TNF-α treatment and reveal a time-dependent reactive oxygen species (ROS) response. The observed decline in ROS levels suggests cellular adaptation to TNF-α, consistent with the literature (30); in this study, such temporal changes in ROS formation inform the analysis of rRNA damage described below.

**Figure 2.**
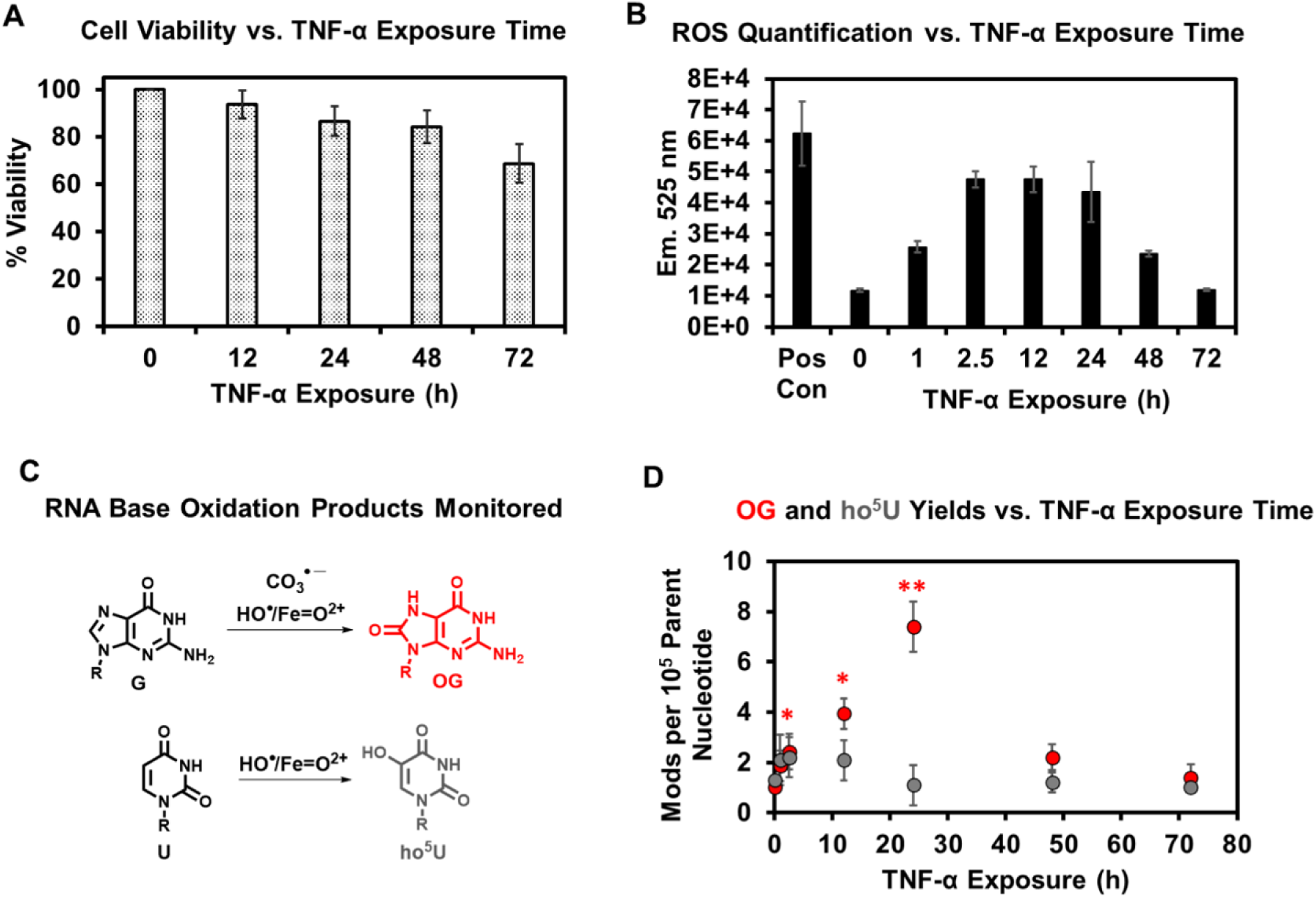
Time-dependent TNF-α exposure in HEK293T cells elicits dynamic RONS production and oxidative modification of total RNA. (A) Cell viability as a function of TNF-α exposure time, determined by a trypan blue exclusion assay. (B) Time-course of oxidative stress measured by DCFH-DA fluorescence during TNF-α treatment. The positive control was 25 μM menadione for 1 h in DMEM with 5% CO_2_ present. (C) Structures of G, OG, U, and ho^5^U, along with the oxygen radicals leading to their formation. The nucleosides OG and ho^5^U are more redox active than their canonical parent nucleosides, which enables their selective quantification by HPLC-UV-ECD (7). (D) The levels of OG and ho^5^U released from total RNA after nuclease/phosphatase digestion, plotted as a function of TNF-α exposure time. The statistical significance was determined by a Student’s *t*-test (\**P* < 0.05 and ** *P* < 0.01).

Fluorescence indicators such as DCFH-DA can reveal an oxidizing cellular environment, but do not identify the specific ROS responsible (31). Therefore, we employed HPLC coupled to UV and electrochemical detectors (HPLC-UV-ECD) to quantify two oxidatively derived lesions in total cellular RNA. The oxidatively modified products OG and ho^5^U were monitored because both remain redox active for electrochemical detection (Figure 2C) (7). The levels of OG report on the generation of CO_3_^•-^ or HO^•^, whereas ho^5^U arises exclusively from HO^•^/Fe=O^2+^ (7). Before TNF-α treatment, OG and ho^5^U levels in total RNA from the cells occurred at ∼1 in 10^5^ of their respective parent nucleotides (Figure 2D). The OG and ho^5^U levels agree well with previously reported values in mammalian cells measured by LC-MS/MS (3, 32), supporting the accuracy of our measurements. Notably, ho^5^U is also enzymatically incorporated into tRNA, contributing to its elevated baseline level.

Following TNF-α exposure for 2.5–24 h, the OG levels rose significantly above baseline, peaking at 7 per 10^5^ G nucleotides at 24 h, before returning to near background at 48 and 72 h (Figures 2D, red; Table S1). In contrast, ho^5^U levels did not increase significantly above baseline (Figure 2D, gray; Table S1). These results implicate CO_3_^•-^ as the RONS responsible for both the DCFH-DA fluorescence signal enhancement (Figure 2B) and oxidation of RNA during inflammation. Together, the DCFH-DA and OG data indicate that RONS production is highest during the early phase of inflammation and declines over time.

### RNA direct nanopore sequencing for rRNA damage during inflammation

RNA direct nanopore sequencing enables the sequencing of RONS-derived lesions in RNA strands. Previous nanopore DNA sequencing studies demonstrated that dOG increases base-calling error rates, facilitating its sequencing (Figure 1A) (33, 34). By analogy, G oxidative lesions in RNA should similarly increase base-calling errors at G sites, and the more structurally distorted hydantoins are expected to induce even greater base miscall rates than OG. Cytosolic and mitochondrial rRNAs from TNF-α–stressed HEK293T cells were sequenced directly, and base-calling error profiles for all G nucleotides were extracted from the reference-aligned reads. The G positions within the 5-nt k-mer of a known epitranscriptomic modification in the rRNA were removed from the analysis because they inherently have increased base-calling errors and would yield false positives (Table S2).

The base-calling error for each G reference nucleotide was calculated and used to generate a population distribution, which was visualized as a 95% confidence interval (CI) plot of G error versus TNF-α exposure time (Figure 3A; top, cytosolic rRNA; bottom, mitochondrial rRNA; see Figure S1 for violin plots). At 0 h, average G base-calling errors were 5.5% for cytosolic rRNA and 3.5% for mitochondrial rRNA. These baseline values reflect two inherent features of the system: (1) RNA direct nanopore sequencing has a high background error rate, and (2) cytosolic rRNA exhibits a higher G error rate than mitochondrial rRNA. The latter likely reflects the greater G content of cytosolic rRNA (34%) compared to mitochondrial rRNA (18%), and the presence of G-rich expansion segments in cytosolic 28S rRNA, which contain G runs that are particularly error-prone in nanopore sequencing (35). Following TNF-α treatment, G base-calling errors at 1 h were indistinguishable from background in both cytosolic and mitochondrial rRNAs (Figure 3A), consistent with OG levels measured by HPLC-UV-ECD at the same time points (Figure 2D). At 12 and 24 h, cytosolic rRNA exhibited significantly elevated G base-calling errors (7.5% on average), whereas mitochondrial rRNA showed a modest increase at 24 h of cytokine exposure (Figure 3A). At 48 h and 72 h, G error rates for both cytosolic and mitochondrial rRNAs returned to near-background levels. These results indicate that TNF-α– induced G oxidation is time-dependent and that the cellular compartment in which rRNA resides influences the extent of oxidative G damage during inflammation.

**Figure 3.**
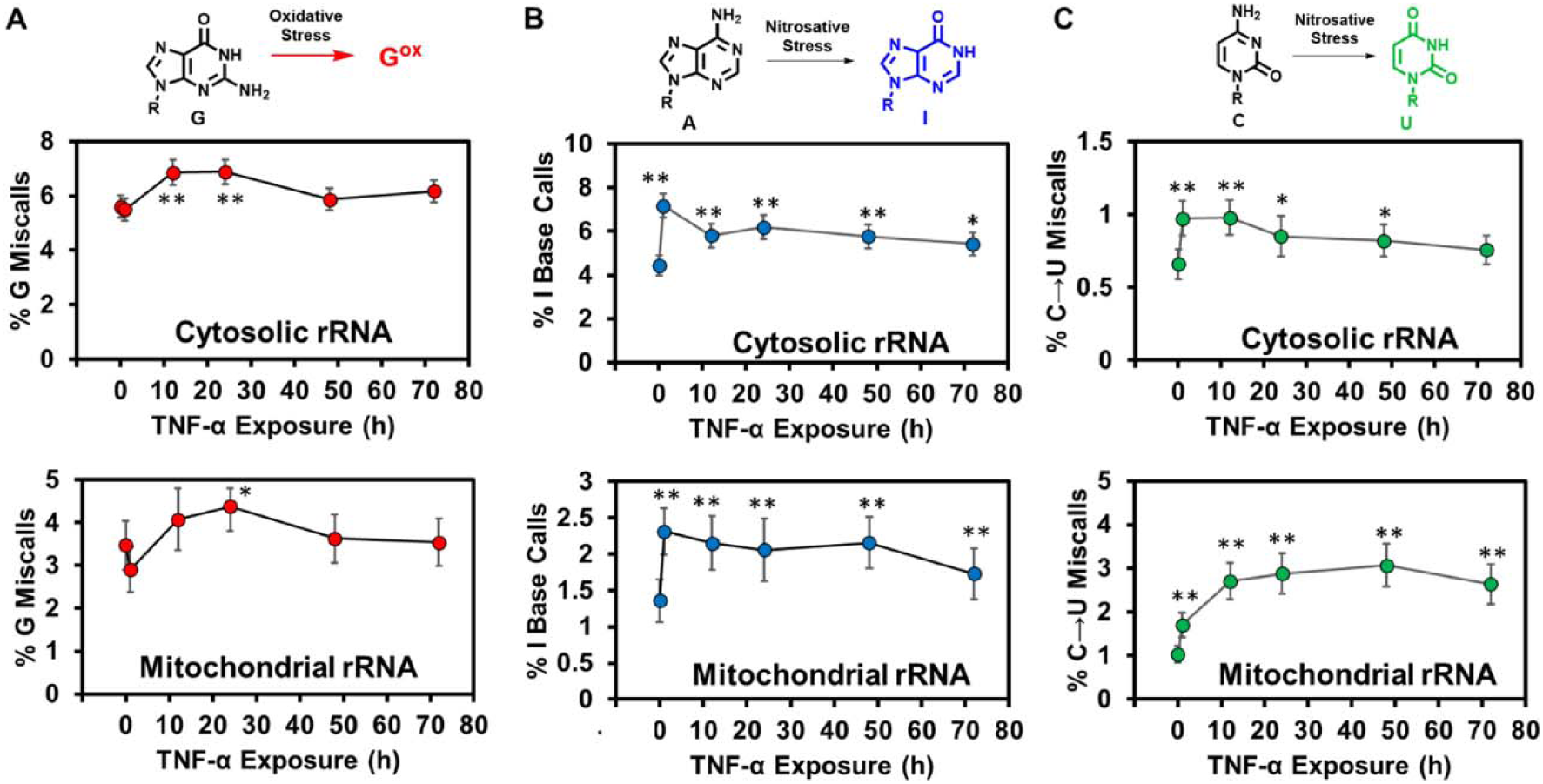
RNA direct nanopore sequencing for rRNAs in cells experiencing inflammatory stress reveals G oxidative modifications and deamination of the A and C heterocycles. (A) Monitoring the time-dependent nanopore base call error at G nucleotides in cytosolic and mitochondrial rRNAs in HEK293T cells exposed to TNF-α at 25 ng/mL. The time dependency in (B) A-to-I deamination and (C) C-to-U deamination in cellular rRNAs analyzed by nanopore sequencing. The Student’s t-test provided significance testing with * *P* < 0.05, ** *P* < 0.01, and *** *P* < 0.001.

Nitrosating agents generated during inflammation induce A-to-I deamination (Figure 1B). The recent nanopore base-calling algorithm Dorado includes a model for direct I identification; we used this tool to quantify I base-call frequency at reference A nucleotides in cytosolic and mitochondrial rRNAs. Sites within a 5-nt k-mer of known epitranscriptomic modifications were excluded from the analyses (Table S2). The I base-call frequencies were displayed as 95% CI plots against TNF-α exposure time, revealing baseline average values of 7% in cytosolic rRNA and 1.4% in mitochondrial rRNA (Figure 3B). One hour after TNF-α treatment, I levels were significantly above background in the rRNA in both compartments (cytosol +2.5%; mitochondria +1%; Figure 3B) and remained elevated throughout the exposure. These results indicate that A-to-I deamination is time-dependent, peaks around 24 h after the initial TNF-α exposure and is sustained in rRNA during prolonged inflammation.

Nitrosating agents formed during inflammation can also induce C-to-U deamination (Figure 1C). Because U is a canonical nucleotide, detecting this reaction by LC-MS/MS is challenging; however, RNA direct nanopore sequencing enables detection of reference C nucleotides miscalled as U to report on the reaction. Aligned sequencing reads were analyzed for U miscall frequency at all C sites, excluding those within the 5-nt k-mer of known epitranscriptomic modifications (Table S2). Before TNF-α treatment, the average C→U miscall frequencies were 0.7% for cytosolic rRNAs and 1.0% for mitochondrial rRNAs (Figure 3C). At 1 h, these values rose significantly to 1.0% and 1.3%, respectively. Frequencies in both compartments remained significantly above background through 48 h. At 72 h, cytosolic rRNAs showed a non-significant increase, whereas mitochondrial rRNAs remained significantly above the background. These findings demonstrate that nanopore sequencing can track inflammation-derived rRNA lesions, revealing distinct temporal patterns for oxidative versus deamination damage, with cellular compartmentalization further influencing the RNA reactivity toward nitrosating reagents.

### RNA direct nanopore sequencing for rRNA damage during oxidative stress

Assessment of rRNA damage from oxidative stress, independent of nitrosating agents, was performed by bolus addition of 100–500 μM H_2_O_2_ directly to HEK293T cells. Oxidation was carried out in PBS supplemented with 20 mM NaHCO_3_. Although these high micromolar concentrations of H_2_O_2_ are supraphysiological, prior studies using intracellular H_2_O_2_ probes have shown that ∼0.1% of the added oxidant enters cells following bolus addition (36). Thus, intracellular H_2_O_2_ concentrations likely increased into the hundreds of nanomolar range, comparable to levels observed in cells undergoing oxidative stress (37).

Titration of H_2_O_2_ into HEK293T cells, followed by RNA direct nanopore sequencing, revealed that miscalls at reference G positions in cytosolic rRNAs increased as a function of the H_2_O_2_ concentration (Figure 4A). A similar trend was observed for mitochondrial rRNAs (Figure 4A). In contrast, no increases in I calls at reference A positions or U calls at reference C positions were detected (Figures 4B and 4C), consistent with the ability of H_2_O_2_ to drive oxidation reactions but not nitrosative reactions. Together, these results demonstrate that oxidative stress promotes G oxidation without deamination, whereas inflammation induces both oxidation and deamination (Figures 3 and 4). These distinct rRNA modification signatures provide a molecular basis to distinguish oxidative stress from inflammation when the source of cellular stress is unclear.

**Figure 4.**
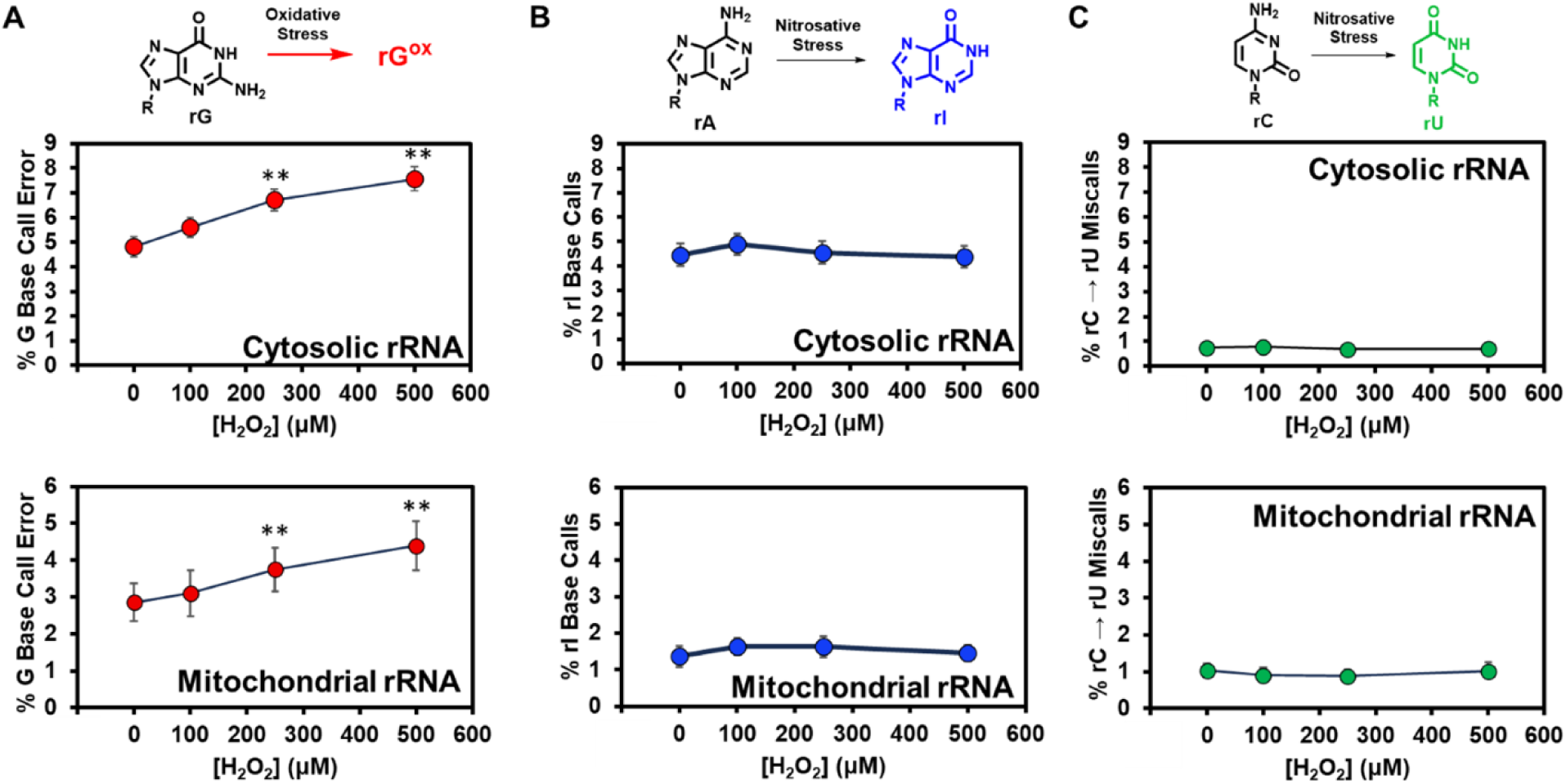
RNA direct nanopore sequencing for rRNAs in cells experiencing oxidative stress reveals only G oxidative modifications. (A) Monitoring the concentration-dependent nanopore base call error at G nucleotides in cytosolic and mitochondrial rRNAs in HEK293T cells exposed to 100-500 μM H_2_O_2_. The concentration dependency in (B) A-to-I deamination and (C) C-to-U deamination in cellular rRNAs analyzed by nanopore sequencing. The Student’s t-test provided significance testing with * *P* < 0.05, ** *P* < 0.01, and *** *P* < 0.001.

### Where do the reactions occur in cytosolic rRNAs?

Further analysis of the RNA direct nanopore sequencing data enables the identification of the specific locations where rRNA undergoes chemical modification within cells. We first focused our inspection on cytosolic rRNAs, as they reacted under both stress conditions (Figure S1). Using the 24 h TNF-α treatment data, we analyzed the 18S and 28S rRNAs individually to determine whether one subunit was more susceptible to damage. Strand-specific analysis revealed that the large subunit rRNA (28S) exhibited a greater increase in G miscalls compared to the small subunit rRNA (18S; Figure S3). This result is consistent with prior findings in *E. coli*, where the large subunit rRNA was more susceptible to H_2_O_2_-induced damage than the small subunit rRNA (38), an observation consistent with its higher G content (Figure S2).

Localization of damaged sites was first established by demonstrating the reproducibility of G miscall signatures. Replicates from the 24-h TNF-α exposure experiment showed strong concordance with scatter plot values fitting the line *x = y* and yielding an *r^2^* = 0.97 (Figure 5A). This confirms that the analyses are reproducible and that, although RNA direct nanopore sequencing is error-prone, the errors are systematic; thus, deviations observed before and after stress treatment cannot be attributed simply to false positives. A scatter plot comparing average G miscalls at 0 h versus 24 h revealed a systematic shift toward higher G miscall frequencies after TNF-α exposure (Figure 5B), as expected. Mapping G miscall differences along the 28S rRNA sequence highlighted regions that we interpret as sites of G oxidation (Figure 5C, red/top). The 15 expansion sequences in the large-subunit rRNA were also plotted (Figure 5C, gray regions), including ES7 and ES27, which are known as “tentacles” due to their extended length and surface exposure (35, 39). The largest increases in G miscalls were observed within these tentacle regions, identifying them as favorable sites of G oxidation. Notably, the tentacles exhibit >80% GC content, resulting in a pattern consistent with the notion that oxidation by CO_3_^•-^ would preferentially target G-rich regions that are surface exposed to react with the diffusible one-electron oxidant (11). In contrast, the peptidyl transferase center (PTC) is not surface exposed nor G-rich and was not found to be preferentially modified (Figure 5C, yellow). However, this stands in contrast with the prior oxidative stress studies in *E. coli* that reported the PTC as a major site of G oxidation (38). Apparently both structure and nucleotide composition matter—the human cytosolic ribosome is much larger than the prokaryotic ribosome, possesses G-rich tentacles that redirect the reaction away from the functionally relevant PTC region of the structure, and is slightly more G-rich (human = 34% and *E. coli* = 31%; Figure S2).

**Figure 5.**
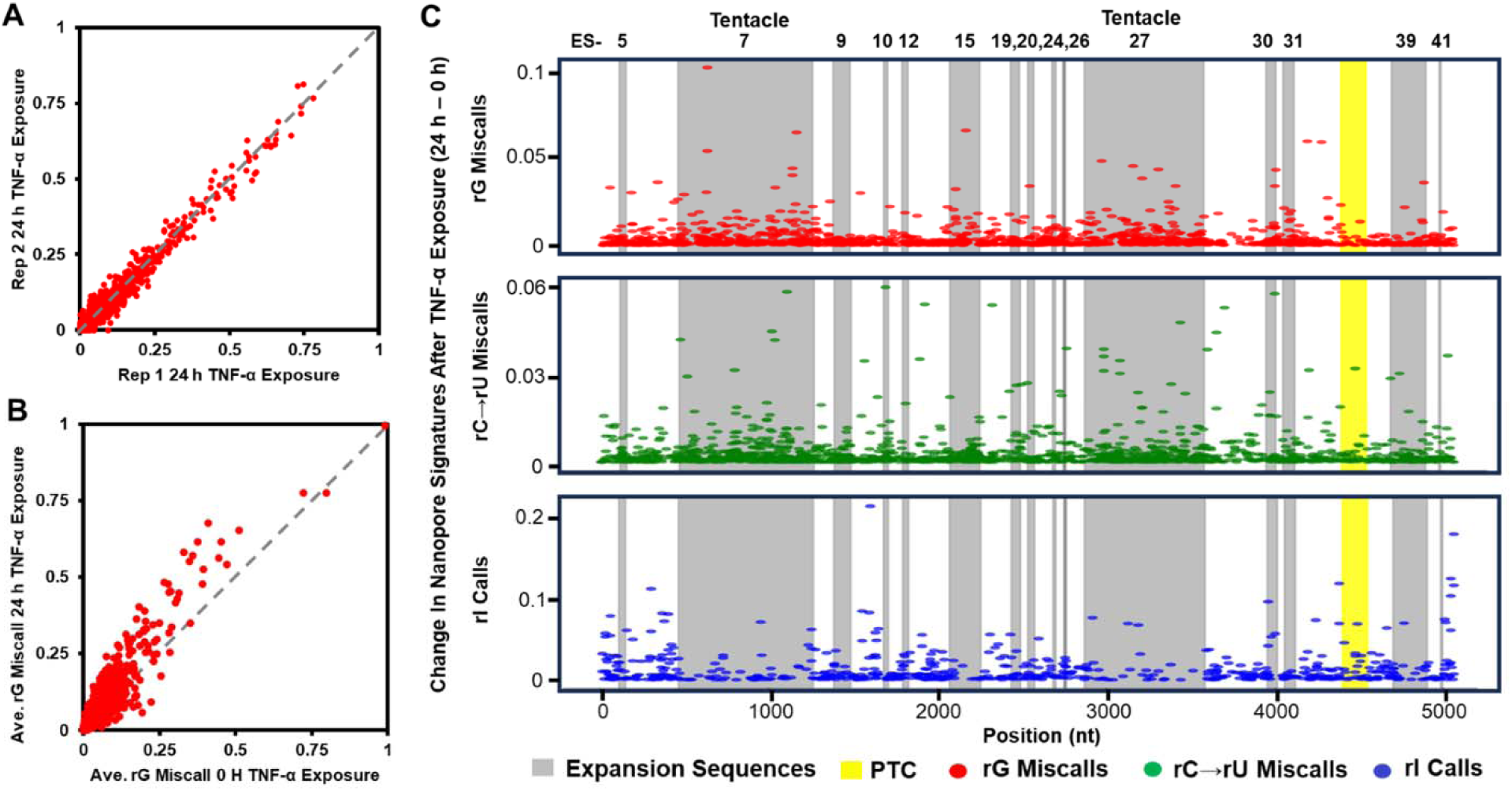
Sites of lesion formation in the 28S rRNA after a 24-h exposure to TNF-α for the generation of inflammatory stress. (A) Scatter plot to demonstrate concordance of the G miscall datasets after a 24-h TNF-α exposure. (B) Scatter plot analysis to determine the G miscall error systematically increases in the 28S rRNA sequenced data after TNF-α exposure. (C) The location of the change in G miscall (red), C-to-U miscall (green), and I calls (blue) versus position in the 28S rRNA to visualize favorable locations for lesions in the rRNA.

Inspection of A and C deamination sites (Figure 5C, blue and green) in the 28S rRNA revealed that C-to-U lesions were enriched within the tentacle regions, whereas A-to-I lesions showed no bias to these regions. The enrichment of C lesions in tentacles is consistent with their C-rich composition and surface exposure, which promote reaction with nitrosating species. In contrast, these same regions are depleted in A nucleotides, explaining why A deamination sites are distributed outside the tentacles. Again, these findings highlight how both nucleotide composition and structural accessibility govern the spatial patterning of rRNA damage under inflammatory stress.

The top 30 most altered G, C, and A sites were mapped onto a cryo-EM structure of the human ribosome (Figure S4) (40). However, tentacles on the 28S rRNA strand have evaded structural inspection due to their surface exposure and dynamic nature; thus, the sites were placed on a 2D map of the 28S rRNA (Figures 6A, 6B red = G-to-G^ox^, 6C blue = A-to-I, and 6D green = C-to-U). Expansion sequences (ES) 7, 15, 27, 30, and 39 (Figure 6A labeled and coded gray) are characterized as having tentacle or tentacle-like properties by projecting from the surface of the structure and having variable sequence compositions, unlike the catalytic core of the structure (35, 41–44). These maps illustrate that the G and C reaction sites preferentially exist in tentacle regions (G^ox^ 53% and C-to-U = 70%; Figures 6B and 6C), and the remaining sites favorably exist in RNA positions near the surface of the structure, consistent with diffusible reactive species preferentially targeting nucleotides that are least sterically shielded. The hairpin structure for the right arm of tentacle ES7 is shown in Figure 6E, showing the GC richness of these sequences. The A-to-I sites reside on the surface of the 28S rRNA and are not enriched in tentacles as a consequence of their depletion in A nucleotides (e.g., ES7; Figure 6E). Closer inspection of the structure around the reactive sites revealed that A-to-I deamination sites generally occur in hairpin loops or large bulges that enhance accessibility (Figure S5). Crucially, these A nucleotides cannot be ADAR substrates, which would require A to be positioned within duplex RNA (27). This structural incompatibility rules out ADAR activity and instead points directly to chemical deamination as the source of the observed I lesions. Enzymatic C-to-U editing can occur, but at a much lower frequency than A-to-I editing, and the results so far support that mRNA is the target of the C deaminases (45). Thus, the C-to-U deamination we see in rRNA is likely a lesion from nitrosative chemistry rather than from enzymatic modification with biological function. Consistently, inspection of the structures of the reactive C nucleotides shows they reside in loops or bulges (Figure S6), where accessibility to nitrosating agents is greatest.

**Figure 6.**
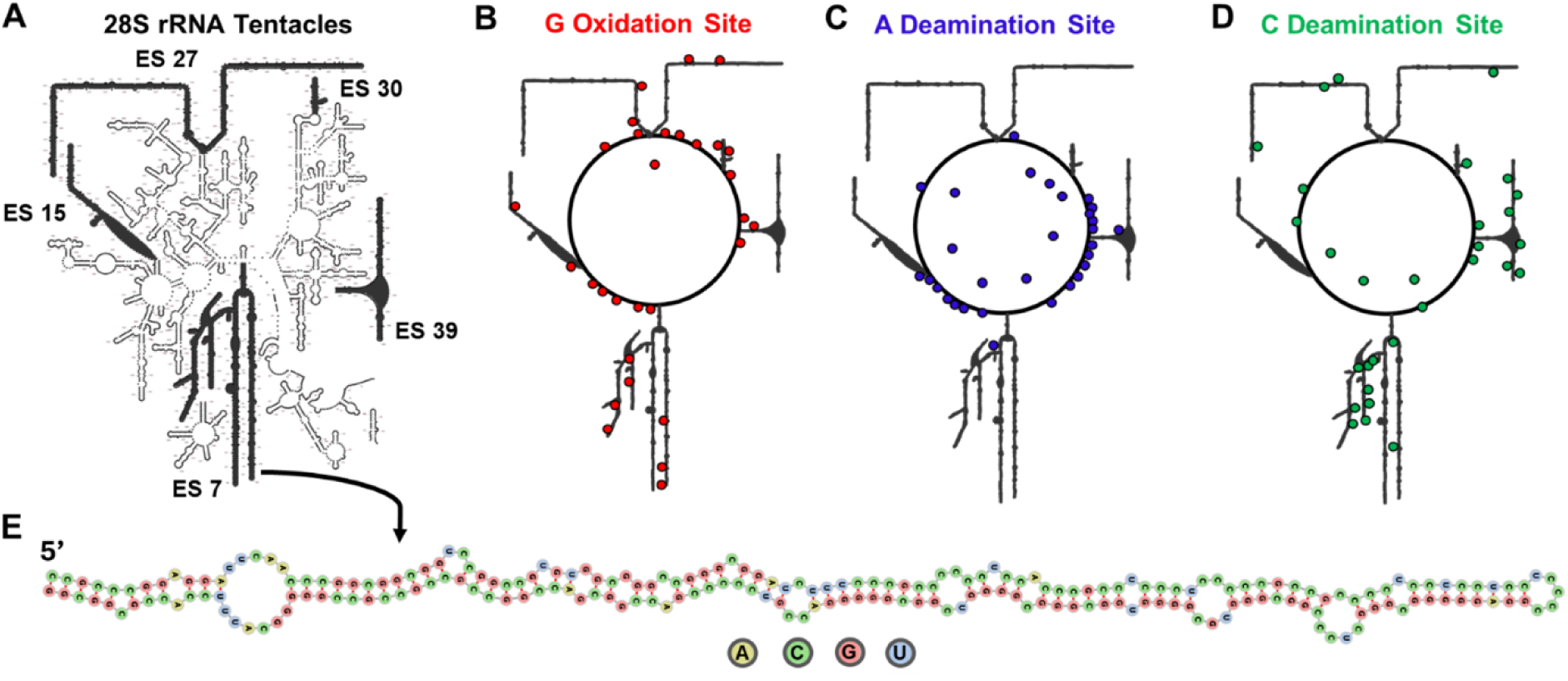
The 28S rRNA tentacles are sites of reaction with inflammation-derived free radicals. (A) A 2D map of the 28S rRNA. Visualization of (B) G oxidation sites, (C) A-to-I deamination sites, and (D) C-to-U deamination sites to illustrate the greater reactivity of the tentacles for G oxidation and C deamination. (E) The secondary structure and sequence for the ES7 tentacle arm showing the G/C richness of the sequences.

Analysis of the 18S, 5.8S, and 5S rRNAs in cytosolic ribosomes revealed that oxidation and deamination events were most pronounced at reactive sites located on the ribosomal surface (Figure S3), reinforcing our findings with the 28S rRNA (Figure 5). Consistent with established mechanisms, H_2_O_2_ treatment produced only increases in rG miscalls, reflecting its specificity for oxidation of the guanine heterocycle. Notably, these results further support that within the 28S rRNA, the tentacle regions represent preferential sites of oxidative damage. Taken together, these findings establish structural exposure, particularly within rRNA tentacles, as a unifying determinant of where chemical damage accumulates across the ribosome.

### Where do the reactions occur in mitochondrial rRNAs?

Structurally, the mitochondrial ribosome represents an ancestral relic, a minimalist biomachine compared to the cytosolic ribosome, while retaining the core functional centers required for protein synthesis. Analysis of G oxidation, C-to-U, and A-to-I lesions in the small (mt-12S) and large (mt-16S) subunit rRNAs revealed reactivity patterns distinct from those observed in cytosolic ribosomal rRNAs (Figure 7A and 7B). In the mt-12S rRNA, all three lesion types appeared broadly dispersed with a few sites showing pronounced increases; a similar pattern was observed in the mt-16S rRNA.

**Figure 7.**
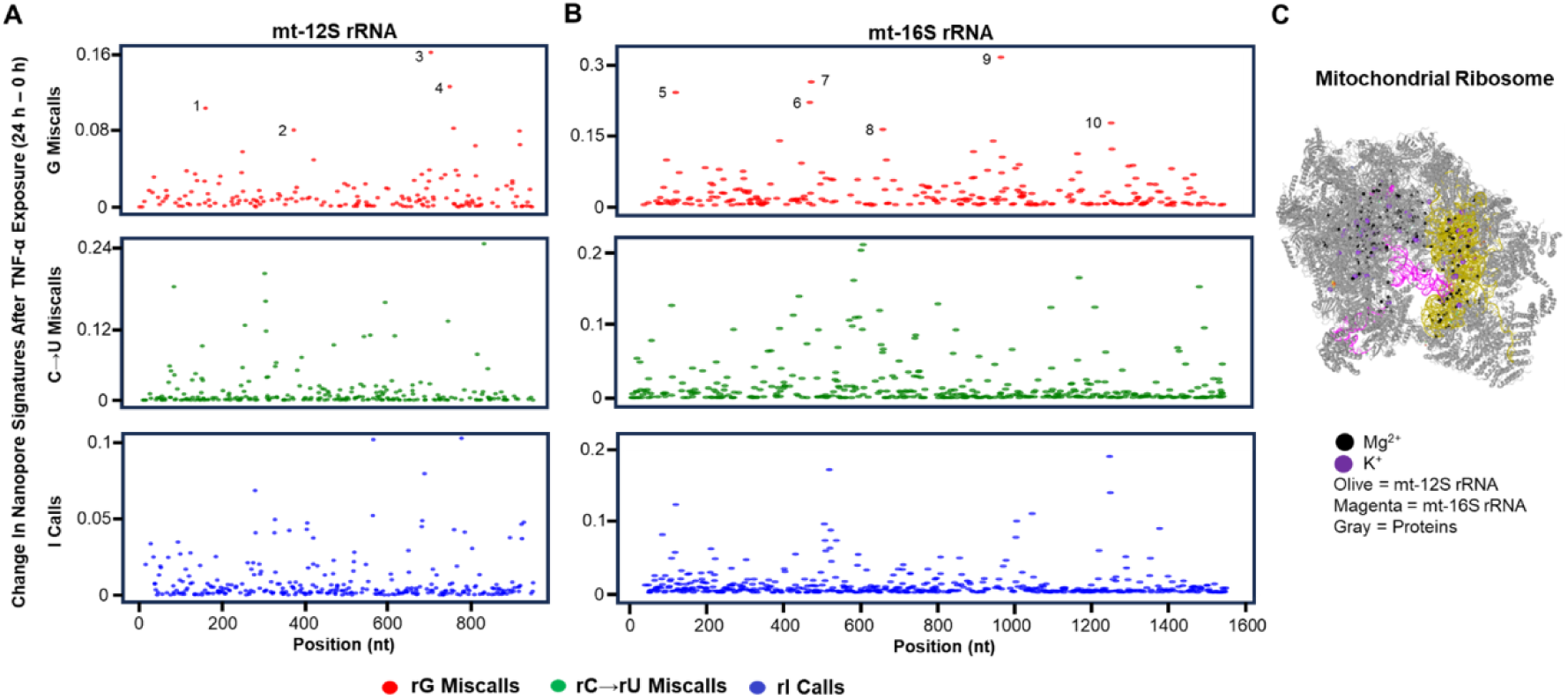
RONS induced lesion patterns in the human mitochondrial ribosome. (A) The G (red, top), C-to-U (green, middle), and A-to-I (blue, bottom) lesion profile for the mt-12S rRNA. (B) The G (red, top), C-to-U (green, middle), and A-to-I (blue, bottom) lesion profile for the mt-16S rRNA. (C) A structure of the mitochondrial ribosome to visualize the mt-12S and mt-16S rRNAs, the Mg^2+^, and K^+^ ions in the structure (pdb 7QI4) (46).

To further investigate, the top 10 candidate rG oxidation sites were identified and numbered in Figures 7A and 7B and mapped onto the 3D ribosome structure. These positions were evaluated for their proximity to Mg^2+^ ions, which in cells can potentially exchange for labile Fe^2+^, a phenomenon that has been demonstrated in *E. coli* ribosomes (47). Such an exchange would increase the prevalence of bicarbonate-Fenton reactions in proximity to the transition metal center. The average distance between reactive G sites and exchange sites was 7.5 Å (range: 4.8–10.0 Å; Table S2), consistent with Fe^2+^-mediated catalysis of oxidation. Supporting this interpretation, studies in yeast showed that H_2_O_2_ treatment under bicarbonate-free conditions (classical Fenton chemistry yielding HO^•^/Fe=O^2+^) produced strand breaks preferentially within ∼7 Å of an exchangeable Mg^2+^/Fe^2+^ binding site in ribosomes (48), paralleling our present observations. Together, these findings suggest that sites of Mg^2+^ binding represent structural hot spots where Fe^2+^ substitution could localize oxidative damage within mitochondrial rRNAs, which may also apply to the cytosolic ribosome reaction sites.

Regarding nitrosative damage, the most reactive sites were dispersed throughout the mitochondrial rRNA sequences (Figures 7A and 7B; green = C-to-U, blue = A-to-I). Mapping the top 10 sites for each deamination type onto the mitochondrial ribosome revealed that they preferentially reside within a structural groove containing functionally critical regions, which are not shielded by the exterior shell of ribosomal proteins (Figure S7). Comparison with prior work in *E. coli* showed a similar pattern of lesions clustering in functional centers of its ribosome, such as the PTC (38). These findings support the conclusion that lesions in both mitochondrial and prokaryotic ribosomes can compromise the fundamental activity of the ribosome to synthesize proteins. RONS targeting to damage nucleotides in the essential functional core of the ribosome thereby provides a direct molecular link between oxidative and inflammatory stress and impaired mitochondrial translation and energy metabolism.

## Discussion

RNA direct nanopore sequencing enables detection of G oxidation through base-calling error analysis and the detection of deamination products inosine and uridine by direct base calling. For G oxidation, this approach identifies reactive G nucleotides but does not reveal the chemical identity of the oxidation products. Nevertheless, the global amount of the presumed dominant oxidation product OG could be measured by HPLC-ECD analysis. Monitoring the most abundant cellular RNA, rRNA, by nanopore analysis showed that during inflammation, cytosolic rRNAs exhibited a ∼1 h delay before G oxidation rose above background, a signal that persisted for 24 h and returned to near-background levels by 48–72 h (Figure 3). In contrast, mitochondrial rRNAs reacted to give a modest increase in G oxidation at 24 h that returned to background at 72 h (Figure 3). Under oxidative stress induced by H_2_O_2_, G oxidation detected in the nanopore data in all rRNAs was dose dependent (Figure 4), with the largest increase observed in cytosolic rRNAs. We hypothesize that the lower reactivity of mitochondrial rRNA reflects two factors: (1) the protective effect of higher HCO_3_^-^concentrations in the mitochondrial matrix (∼100 mM vs. ∼20 mM in the cytosol) maintained by the alkaline pH and continuous CO_2_ generation through the citric acid cycle (7, 24), and (2) the reduced G content of mitochondrial rRNAs (18% vs. 34% in cytosolic rRNAs; Figure S2) along with the absence of G-rich ribosomal expansion sequences that protrude from the ribosome surface and are more exposed to diffusible reactive species. These findings suggest that both the mode of generation of oxidative stress and the compartmentalization of RNA substrates define when and where endogenous oxidative stress induces G oxidation.

Evaluation of A and C deamination in response to nitrosating agents provided additional insights into the endogenous chemistry that introduces lesions in rRNA. RNA direct nanopore sequencing can detect the deamination product I using the modification-aware base caller and U using the standard base caller. During oxidative stress induced by H_2_O_2_, deamination of A or C was not observed, consistent with the underlying reaction mechanism (Figure 4). In contrast, when TNF-α was used to induce inflammation, deamination lesions appeared earlier than G oxidation in both cytosolic and mitochondrial rRNAs (Figure 3). This timing suggests that nitric oxide synthases are induced before superoxide-generating oxidoreductases, with the early burst of ^•^NO driving A and C deamination through formation of nitrosating species (25). Furthermore, A and C deamination persisted throughout TNF-α exposure, whereas G oxidation peaked at 12–24 h post-treatment. These findings imply distinct induction kinetics for ^•^NO-versus O_2_^•-^-generating enzymes, consistent with prior reports (49, 50).

An additional observation regarding rRNA lesions is that G oxidation displays dependency on the cellular compartment in which the RNA resides during inflammation, whereas A and C deamination reactions are not strongly compartment dependent. The present findings are interesting because prior in vitro studies found that HCO_3_^-^inhibits N_2_O_3_-mediated deamination reactions (51), which should impose compartment dependency on rRNA deamination similar to the oxidation reactions. However, the cellular context of these reactions resolves this apparent contradiction. Nitric oxide, ^•^NO_2_, and O_2_ are hydrophobic and concentrate in lipid bilayers, thereby promoting N_2_O_3_ generation at membranes (25). Mitochondrial ribosomes, which exclusively translate membrane-associated electron transport chain proteins, are anchored to the inner mitochondrial membrane to deliver the newly formed protein (52). Similarly, a large fraction of cytosolic ribosomes is membrane-associated with the endoplasmic reticulum (53). Thus, membrane localization of ribosomes creates a favorable environment where N_2_O_3_ formation is elevated near the rRNA that is the target of the deamination reaction. Thus, HCO_3_^-^in the bulk solution is expect to have minimal impact on the chemistry occurring near membrane surfaces. This feature of N_2_O_3_ formation in cells assists in understanding the compartment-independent deamination of A and C observed in cellular rRNAs.

The cytosolic ribosomal RNA expansion sequences containing GC-rich tentacles were particularly susceptible to G oxidation and C deamination. Notably, the evolution of the eukaryotic ribosome from *E. coli* to humans has increased the mass of the large subunit rRNA by more than 40%, with most of the added sequences being unusually GC-rich (>80%) and exhibiting pronounced GC skew (Figure 6E) (54, 55). No crystallographic or cryo-EM structures are available for these tentacles, likely because they extend into solution and are too dynamic to pack into a regular array. Expansion sequences have been ascribed roles in translation fidelity and as platforms for initiation and biogenesis factors (54). The paucity of structural information for these regions continues to inspire investigation of the expansion sequences in the human ribosome. The G-rich stretches of these sequences have been proposed to fold into G-quadruplexes (see Figure 6E for an example) (35); however, their nearly complementary C-rich regions cannot form stable i-motifs because such structures are disfavored by the furanose conformation in RNA (56), leaving long poly-C tracts structurally orphaned. In this work, we found ∼55% of G oxidation sites and 70% of C deamination sites were located in expansion sequences. Both modifications are expected to alter structure, either by destabilizing G:C base pairs or by introducing new G:U wobble pairs. Functionally, the GC-rich tentacles appear to act as preferential targets for free radicals generated during oxidative stress (e.g., CO_3_^•−^) or nitrosative stress, where ^•^NO and ^•^NO_2_ combine to form the potent nitrosating agent N_2_O_3_. These observations suggest that human cytosolic ribosome architecture may have evolved to localize radical damage to peripheral, non-catalytic regions, thereby protecting essential functional centers and providing cytosolic ribosomes with longer lifetimes.

## Materials and Methods

### Cell culture experiments

The HEK293T cells were grown in DMEM medium supplemented with 10% FBS, 1x GlutaMAX™, 1x non-essential amino acids, and 25 μg/mL gentamicin. The cells were grown in a cell culture incubator under a blanket of 5% CO_2_ with 80% humidity at 37 °C. When they reached ∼50% confluency, 10^6^ or 10^7^ cells were treated with 25 ng/mL of TNF-α for 1-72 h in sterile cell culture flasks. For cells exposed to TNF-α for >24 h, the medium and TNF-α were replaced every 24 h. The cells were harvested by centrifugation at each time point of the study. Oxidative stress by bolus addition of H_2_O_2_ was conducted on 10^6^ cells harvested from standard culture conditions in DMEM medium. The cells were pelleted by centrifugation, washed with 1x PBS, and then resuspended in 1X PBS with 20 mM cell-grade NaHCO_3_. After incubating the cells for 30 min, a bolus of 100, 250, or 500 μM H_2_O_2_ was added, and they were allowed to react for 15 min before quenching the reaction by centrifugation and decanting the medium. All cell pellets were stored at -80 °C until studied.

### Nanopore sequencing experiments

Total RNA was extracted from the cells using the Zymogen RNA extraction kit following the manufacturer’s protocol, with one exception. To the extraction buffers was added 100 μM desferrioxamine, an iron chelator, and 100 μM butylated hydroxytoluene, an antioxidant, to minimize unwanted RNA oxidation during the extraction process, which is a known issue (57). RNA integrity was determined by agarose gel electrophoresis. Before library preparation, 2 μg of total RNA was 3′ poly-A tailed using a commercial poly-A tailing kit following the instructions (LGC Biosearch Technologies). Library preparation was conducted using the SQK-RNA004 kit (ONT) following the manufacturer’s protocol. The library-prepared RNA was sequenced using the RNA flow cell from ONT running default parameters that pass reads with Q > 8. The passed reads were base called with Dorado in fast mode with default parameters, or in hac mode with the inosine-aware base call model (https://github.com/nanoporetech/dorado). The data were analyzed using IGV,(58) Excel, Origin, bash, and Python using the pysam, pandas, matplotlib, and numpy libraries. Data for the plots were filtered to remove positions within a 5-nt k-mer of a known rRNA epitranscriptomic modification and the read depth was > 100. All sequencing data and in-house code are publicly available.

## Acknowledgments

This work was supported by the National Institute of General Medical Sciences via grant no. R35 GM145237. The computational resources used were partially funded by the NIH Shared Instrumentation Grant 1S10OD021644-01A1. **Author Contributions.** Conceptualization and methodology: A.M.F. and C.J.B. Cell culture: A.M.F. Nanopore Sequencing and Data Analysis: A.M.F. HPLC-UV-ECD: J.C.D. Writing: A.M.F. and C.J.B. Supervision and Funding Acquisition: C.J.B. **Competing Interests.** A.M.F. and C.J.B. have a patent licensed to Electronic BioSciences, and A.M.F. occasionally consults on nucleic acid chemistry for Electronic BioSciences. **Data and Materials Availability.** The base called data for all sequencing experiments, the nanopore analysis data, and the Python code for the analysis have been publicly deposited at Zenodo, a public repository for data at https://doi.org/10.5281/zenodo.13255917, and https://doi.org/10.5281/zenodo.17095368 which are searchable in the OpenAIRE explorer.

## Supporting Information

Complete Materials and Methods

Study Limitations

Figures S1-S8

Tables S1-S4

