## Supplemental Information for "Ribosomal RNA tentacles are targets of free radical damage in mammalian cells during oxidative and inflammatory stress"

**Item	Page**

**Materials and Methods**	S2

**Limitations**	S3

**Table S1**. The DCFH-DA values obtained for the ROS measurements during TNF-α exposure. S4

**Table S2**. The HPLC-UV-ECD values obtained for measuring OG and ho^5^U.	S5

**Table S3**. The positions and identities of the human rRNA epitranscriptomic modifications. S6

**Table S4**. Distances between 10 mitochondrial G oxidation sites and the nearest Mg^2+^ ion. S13

**Figure S1**. Violin plots for the same data used to construct the 95% CI plots in Figs. 3 and 4. S14

**Figure S2**. Analysis of changes in G oxidation in the 18S vs. 28S rRNAs.	S16

**Figure S3**. Nucleotide compositions of *E. coli* and human rRNAs.	S17

**Figure S4**. Structure of human cytosolic ribosome with lesions mapped on the structure. S18

**Figure S5**. 2D structures for sites of A-to-I deamination in the cytosolic ribosome.	S19

**Figure S6**. 2D structures for sites of C-to-U deamination in the cytosolic ribosome.	S20

**Figure S7**. Comparative plots for the 5S, 5.8S, and 18S rRNAs before and after 24 h TNF-α exposure.	S21

**Figure S8.** Mitochondrial ribosome with top lesion sites mapped on the structure.	S23

**References**	S24

**Materials and Methods**

**Cell culture experiments.** The HEK293T cells were grown in DMEM medium supplemented with 10% FBS, 1x GlutaMAX™, 1x non-essential amino acids, and 25 μg/mL gentamicin. The cells were grown in a cell culture incubator under a blanket of 5% CO_2_ with 80% humidity at 37 °C. When they reached ~50% confluency, 10^6^ or 10^7^ cells were treated with 25 ng/mL of TNF-α for 1-72 h in sterile cell culture flasks. For cells exposed to TNF-α for >24 h, the medium and TNF-α were replaced every 24 h. The cells were harvested by centrifugation at each time point of the study. Oxidative stress by bolus addition of H_2_O_2_ was conducted on 10^6^ cells harvested from standard culture conditions in DMEM medium. The cells were pelleted by centrifugation, washed with 1x PBS, and then resuspended in 1X PBS with 20 mM cell-grade NaHCO_3_. After incubating the cells for 30 min, a bolus of 100, 250, or 500 μM H_2_O_2_ was added, and they were allowed to react for 15 min before quenching the reaction by centrifugation and decanting the medium. All cell pellets were stored at -80 °C until studied. The DCFH-DA assay (Caymon Chemicals Cat # 601520) was conducted as outlined in the manufacturer’s protocol.

**Nanopore sequencing experiments**. Total RNA was extracted from the cells using the Zymogen RNA extraction kit following the manufacturer’s protocol, with one exception. To the extraction buffers was added 100 μM desferrioxamine, an iron chelator, and 100 μM butylated hydroxy toluene, an antioxidant, to minimize unwanted RNA oxidation during the extraction process, which is a known issue (1). The RNA integrity was determined by agarose gel electrophoresis. Before library preparation, 2 μg of total RNA was 3′ poly-A tailed using a commercial poly-A tailing kit following the instructions (LGC Biosearch Technologies). Library preparation was conducted using the SQK-RNA004 kit (ONT) following the manufacturer’s protocol. The library-prepared RNA was sequenced using the RNA flow cell from ONT running default parameters that pass reads with Q > 8. The passed reads were base called with Dorado in the fast mode with default parameters, or in hac mode with the inosine-aware base call model (<https://github.com/nanoporetech/dorado>). The reads in the BAM file format were aligned using Dorado with default parameters. The G oxidation and C deamination data were obtained from base calling using the fast mode in Dorado, while the inosine-aware data were obtained using the hac mode in Dorado.

The aligned bam files were inspected for changes associated with lesions in the RNA. The G oxidation sites were evaluated as follows. The base calls at each site were obtained from the aligned bam files using in-house Python code running the pysam library to yield the base calls at each position in tsv file format. The base calls at each reference G were selected using grep and then filtered for those that had a read depth of >100 using the awk function in bash. When comparing data sets between no reaction and reaction, the intercept files were obtained using the awk function in bash. The data were plotted in either Origin, Excel, or with Python running pandas, matplotlib, and numpy libraries. The inosine base calls were obtained using the m^6^A-inosine aware base caller to generate base call files in bam format. The base call data were aligned to the reference with Dorado to yield reference-aligned bam files. The bed files with position-dependent m^6^A and I base calls were filtered to first obtain bed files with I data using grep. The I bed files were filtered for those with read depths >100 using the awk function in bash. Lastly, intersection bed files for no reaction vs. those after reaction were again obtained using the awk function in bash. Plotting was achieved as described for the G oxidation sites. The C-to-U deamination sites were obtained following the same approach as the G oxidation analysis, but the data were analyzed for the extent of U miscalls at reference C positions in the rRNAs.

**Limitations.** There are two central limitations to this study. The first concerns the use of nanopore sequencing to identify rare RNA modifications introduced by chemical reactions rather than enzymatic processes. The probability of detecting a modification depends on whether the reaction occurs frequently enough at a given site to produce a signal that rises above the background sequencing error rate. In the case of G oxidation, modifications are biased toward G-rich regions, which in the 28S rRNA are found in G runs within tentacles. This presents a complication because nanopore sequencing exhibits its highest error rates in homopolymer runs, which are the very regions where G oxidation is most likely. However, these sequencing errors are highly reproducible across replicates (Figure 5A), and the oxidation of a G disrupts a homopolymer run, producing a distinct change in the error profile (Figure 5B). It is this reproducible change that we track in G runs.

By targeting rRNA as the oxidation substrate in human cells, the analysis may already start with a relatively high baseline of OG sites (~1 per 10^3^ Gs in *E. coli*) (2). Two additional factors likely enhance the signal above background. (a) If pre-existing OG lesions are located at hotspot positions, their intrinsic lability toward further oxidation will result in site-specific hydantoin product formation (Figure 1A). These hydantoin products likely yield highly error-prone signatures during nanopore base calling, leading to site-specific signal amplification. (b) Under oxidative stress, rRNA likely bears a disproportionate share of the total transcriptome damage, resulting in a higher local density of modified sites than would be evident from bulk measurements (Figure 2D). Together, these features, site selectivity and elevated product yield, contribute to signals that rise above the inherent error rate of the sequencer.

When oxidation occurs within G runs, the resulting signal can only be localized at the level of the k-mer (~5 nt), rather than to a single nucleotide.

These arguments for signal enhancement hold for A-to-I and C-to-U deamination sites. To our advantage the nanopore base caller is calibrated to find these modifications, one having a model for modification aware base calling (I) and the other being native to RNA (U).

Finally, we point out that the data are not quantitative between G oxidation, A deamination, or C deamination because different features and/or base call algorithms were used for the data analysis.

The second limitation of the present study is in regards to growing the HEK293T cells under standard incubator conditions with atmospheric O_2_ present at 20%. The physiological O_2_ levels in humans are tissue dependent and at values around 5%; however, atmospheric O_2_ levels remain standard for modern cell culture incubators. The consequence of the high O_2_ levels is higher oxidative stress and RNA damage. High O_2_ levels can also change how the cell responds during inflammation to form reactive radicals (e.g., NAD(P)H oxidoreductases forming O_2_^•-^ vs. H_2_O_2_). The key point is that the levels of RNA damage in these cell culture assays are elevated above what would likely occur in the tissues of an organism.

**Table S1**. The DCFH-DA values obtained for the ROS measurements during TNF-α exposure.

|  |  | TNF-α Exposure Time (h) | | | | | | |
| --- | --- | --- | --- | --- | --- | --- | --- | --- |
| Treatment | Pos Con | 0 | 1 | 2.5 | 12 | 24 | 48 | 72 |
|  | 7.41E+04 | 1.17E+04 | 2.78E+04 | 4.61E+04 | 4.75E+04 | 3.88E+04 | 2.45E+04 | 1.19E+04 |
|  | 5.84E+04 | 1.13E+04 | 2.52E+04 | 4.61E+04 | 4.34E+04 | 5.48E+04 | 2.34E+04 | 1.23E+04 |
|  | 5.44E+04 | 1.23E+04 | 2.43E+04 | 5.05E+04 | 5.18E+04 | 3.70E+04 | 2.27E+04 | 1.18E+04 |
| Ave | 6.23E+04 | 1.18E+04 | 2.58E+04 | 4.76E+04 | 4.76E+04 | 4.35E+04 | 2.36E+04 | 1.20E+04 |
| Std. Dev. | 1.04E+04 | 4.93E+02 | 1.82E+03 | 2.53E+03 | 4.23E+03 | 9.78E+03 | 9.02E+02 | 2.95E+02 |

Pos Con = positive control (25 μM menadione for 1 h)

**Table S2**. The HPLC-UV-ECD values obtained for measuring OG and ho^5^U.

| OG | x10^5^ |  |  |  |
| --- | --- | --- | --- | --- |
| time (h) | Rep 1 | Rep 2 | Ave | Std. Dev. |
| 0.0 | 1.1 | 1.0 | 1.0 | 0.1 |
| 1.0 | 1.8 | 2.0 | 1.9 | 0.6 |
| 2.5 | 2.8 | 2.1 | 2.4 | 0.7 |
| 12.0 | 3.9 | 3.9 | 3.9 | 0.6 |
| 24.0 | 8.1 | 6.7 | 7.4 | 1.0 |
| 48.0 | 1.9 | 2.5 | 2.2 | 0.5 |
| 72.0 | 1.5 | 1.3 | 1.4 | 0.5 |

| ho^5^U | x10^5^ |  |  |  |
| --- | --- | --- | --- | --- |
| time (h) | Rep 1 | Rep 2 | Ave | Std. Dev. |
| 0.0 | 1.3 | 1.5 | 1.4 | 0.2 |
| 1.0 | 2.8 | 1.5 | 2.1 | 1.0 |
| 2.5 | 2.7 | 1.6 | 2.2 | 0.8 |
| 12.0 | 1.4 | 2.9 | 2.1 | 0.8 |
| 24.0 | 0.5 | 1.7 | 1.1 | 0.8 |
| 48.0 | 0.8 | 1.6 | 1.2 | 0.4 |
| 72.0 | 0.8 | 1.1 | 1.0 | 0.1 |

**Table S3**. The positions and identities of the human rRNA epitranscriptomic modifications.*

| **rRNA** | **Position** | **Modification** |
| --- | --- | --- |
| 5.8S | 14 | Um |
| 5.8S | 55 | Ψ |
| 5.8S | 69 | Ψ |
| 5.8S | 75 | Gm |
| 18S | 27 | Am |
| 18S | 34 | Ψ |
| 18S | 36 | Ψ |
| 18S | 93 | Ψ |
| 18S | 99 | Am |
| 18S | 105 | Ψ |
| 18S | 109 | Ψ |
| 18S | 116 | Um |
| 18S | 119 | Ψ |
| 18S | 121 | Um |
| 18S | 159 | Am |
| 18S | 166 | Am |
| 18S | 172 | Um |
| 18S | 174 | Cm |
| 18S | 210 | Ψ |
| 18S | 218 | Ψ |
| 18S | 296 | Ψ |
| 18S | 300 | Ψ |
| 18S | 354 | Um |
| 18S | 406 | Ψ |
| 18S | 428 | Um |
| 18S | 436 | Gm |
| 18S | 462 | Cm |
| 18S | 468 | Am |
| 18S | 484 | Am |
| 18S | 509 | Gm |
| 18S | 512 | Am |
| 18S | 514 | Ψ |
| 18S | 517 | Cm |
| 18S | 556 | Ψ |
| 18S | 572 | Ψ |
| 18S | 573 | Ψ |
| 18S | 576 | Am |
| 18S | 581 | U? |
| 18S | 590 | Am |
| 18S | 601 | Gm |
| 18S | 609 | Ψ |
| 18S | 621 | Cm |
| 18S | 627 | Um |
| 18S | 644 | Gm |
| 18S | 649 | Ψ |
| 18S | 651 | Ψ |
| 18S | 667 | Ψ |
| 18S | 668 | Am |
| 18S | 681 | Ψ |
| 18S | 683 | Gm |
| 18S | 686 | Ψ |
| 18S | 769 | Ψ |
| 18S | 770 | Ψ |
| 18S | 797 | Cm |
| 18S | 799 | Um |
| 18S | 801 | Ψ |
| 18S | 804 | Ψ |
| 18S | 814 | Ψ |
| 18S | 815 | Ψ |
| 18S | 822 | Ψ |
| 18S | 863 | Ψ |
| 18S | 866 | Ψ |
| 18S | 867 | Gm |
| 18S | 889 | Ψ |
| 18S | 897 | Ψ |
| 18S | 918 | Ψ |
| 18S | 966 | Ψ |
| 18S | 1003 | Ψ |
| 18S | 1004 | Ψ |
| 18S | 1031 | Am |
| 18S | 1045 | Ψ |
| 18S | 1046 | Ψ |
| 18S | 1056 | Ψ |
| 18S | 1061 | Ψ |
| 18S | 1081 | Ψ |
| 18S | 1136 | Ψ |
| 18S | 1174 | Ψ |
| 18S | 1177 | Ψ |
| 18S | 1186 | Ψ |
| 18S | 1219 | C? |
| 18S | 1232 | Ψ |
| 18S | 1238 | Ψ |
| 18S | 1239 | Ψ |
| 18S | 1244 | Ψ |
| 18S | 1248 | m1acp3Ψ |
| 18S | 1268 | C? |
| 18S | 1272 | Cm |
| 18S | 1288 | Um |
| 18S | 1315 | Ψ |
| 18S | 1326 | Um |
| 18S | 1328 | Gm |
| 18S | 1337 | ac4C |
| 18S | 1347 | Ψ |
| 18S | 1359 | Ψ |
| 18S | 1367 | Ψ |
| 18S | 1383 | Am |
| 18S | 1391 | Cm |
| 18S | 1400 | Ψ |
| 18S | 1440 | Cm |
| 18S | 1442 | Um |
| 18S | 1445 | Ψ |
| 18S | 1447 | Gm |
| 18S | 1463 | U? |
| 18S | 1490 | Gm |
| 18S | 1596 | Ψ |
| 18S | 1625 | Ψ |
| 18S | 1639 | m7G |
| 18S | 1643 | Ψ |
| 18S | 1668 | Um |
| 18S | 1678 | Am |
| 18S | 1692 | Ψ |
| 18S | 1703 | Cm |
| 18S | 1804 | Um |
| 18S | 1832 | m6A |
| 18S | 1839 | Ψ |
| 18S | 1842 | ac4C |
| 18S | 1850 | m62A |
| 18S | 1851 | m62A |
| 28S | 224 | Ψ |
| 28S | 398 | Am |
| 28S | 400 | Am |
| 28S | 1316 | Gm |
| 28S | 1322 | m1A |
| 28S | 1323 | Am |
| 28S | 1326 | Am |
| 28S | 1340 | Cm |
| 28S | 1522 | Gm |
| 28S | 1524 | Am |
| 28S | 1534 | Am |
| 28S | 1536 | Ψ |
| 28S | 1582 | Ψ |
| 28S | 1625 | Gm |
| 28S | 1677 | Ψ |
| 28S | 1683 | Ψ |
| 28S | 1700 | U? |
| 28S | 1744 | Ψ |
| 28S | 1760 | Gm |
| 28S | 1773 | Um |
| 28S | 1779 | Ψ |
| 28S | 1781 | Ψ |
| 28S | 1782 | Ψ |
| 28S | 1792 | Ψ |
| 28S | 1859 | m4C |
| 28S | 1860 | Ψ |
| 28S | 1862 | Ψ |
| 28S | 1871 | Am |
| 28S | 1881 | Cm |
| 28S | 2351 | Cm |
| 28S | 2363 | Am |
| 28S | 2364 | Gm |
| 28S | 2365 | Cm |
| 28S | 2401 | Am |
| 28S | 2415 | Um |
| 28S | 2422 | Cm |
| 28S | 2424 | Gm |
| 28S | 2508 | Ψ |
| 28S | 2632 | Ψ |
| 28S | 2787 | Am |
| 28S | 2804 | Cm |
| 28S | 2815 | Am |
| 28S | 2824 | Cm |
| 28S | 2837 | Um |
| 28S | 2839 | Ψ |
| 28S | 2843 | Ψ |
| 28S | 2856 | U? |
| 28S | 2861 | Cm |
| 28S | 2876 | Gm |
| 28S | 3627 | Gm |
| 28S | 3637 | Ψ |
| 28S | 3639 | Ψ |
| 28S | 3669 | Gm |
| 28S | 3695 | Ψ |
| 28S | 3701 | Cm |
| 28S | 3715 | Ψ |
| 28S | 3718 | Am |
| 28S | 3723 | Am |
| 28S | 3724 | Am |
| 28S | 3730 | Ψ |
| 28S | 3734 | Ψ |
| 28S | 3744 | Gm |
| 28S | 3758 | Ψ |
| 28S | 3760 | Am |
| 28S | 3762 | Ψ |
| 28S | 3764 | Ψ |
| 28S | 3768 | Ψ |
| 28S | 3770 | Ψ |
| 28S | 3782 | m5C |
| 28S | 3785 | Am |
| 28S | 3792 | Gm |
| 28S | 3808 | Cm |
| 28S | 3818 | Ψm |
| 28S | 3822 | Ψ |
| 28S | 3825 | Am |
| 28S | 3830 | Am |
| 28S | 3841 | Cm |
| 28S | 3844 | Ψ |
| 28S | 3851 | Ψ |
| 28S | 3853 | Ψ |
| 28S | 3867 | Am |
| 28S | 3869 | Cm |
| 28S | 3884 | Ψ |
| 28S | 3887 | Cm |
| 28S | 3899 | Gm |
| 28S | 3920 | Ψ |
| 28S | 3925 | Um |
| 28S | 3944 | Gm |
| 28S | 3959 | Ψ |
| 28S | 4042 | Gm |
| 28S | 4054 | Cm |
| 28S | 4196 | Gm |
| 28S | 4220 | m6A |
| 28S | 4227 | Um |
| 28S | 4228 | Gm |
| 28S | 4293 | Ψ |
| 28S | 4296 | Ψ |
| 28S | 4299 | Ψ |
| 28S | 4306 | Um |
| 28S | 4312 | Ψ |
| 28S | 4353 | Ψ |
| 28S | 4361 | Ψ |
| 28S | 4370 | Gm |
| 28S | 4392 | Gm |
| 28S | 4403 | Ψ |
| 28S | 4420 | Ψ |
| 28S | 4423 | Ψ |
| 28S | 4431 | Ψ |
| 28S | 4442 | Ψ |
| 28S | 4447 | m5C |
| 28S | 4456 | Cm |
| 28S | 4457 | Ψ |
| 28S | 4471 | Ψ |
| 28S | 4493 | Ψ |
| 28S | 4494 | Gm |
| 28S | 4498 | Um |
| 28S | 4499 | Gm |
| 28S | 4500 | Ψ |
| 28S | 4521 | Ψ |
| 28S | 4523 | Am |
| 28S | 4530 | m3U |
| 28S | 4531 | Ψ |
| 28S | 4532 | Ψ |
| 28S | 4536 | Cm |
| 28S | 4552 | Ψ |
| 28S | 4569 | Ψ |
| 28S | 4571 | Am |
| 28S | 4576 | Ψ |
| 28S | 4579 | Ψ |
| 28S | 4590 | Am |
| 28S | 4599 | U? |
| 28S | 4618 | Gm |
| 28S | 4620 | Um |
| 28S | 4623 | Gm |
| 28S | 4628 | Ψ |
| 28S | 4636 | Ψ |
| 28S | 4637 | Gm |
| 28S | 4673 | Ψ |
| 28S | 4689 | Ψ |
| 28S | 4972 | Ψ |
| 28S | 4973 | Ψ |
| 28S | 5001 | Ψ |
| 28S | 5010 | Ψ |

*The positions and modification identities were compiled from literature resources (3-5). Those with “?” were newly reported modification sites obtained from RNA direct nanopore sequencing data (6), in which the chemical identity of the modification remains to be verified.

**Table S4**. Distances between 10 mitochondrial G oxidation sites and the nearest Mg^2+^ ion.

| rRNA | Position | Distance to Mg^2+^ (Å) |
| --- | --- | --- |
| 16S | 473 | 9.6 |
| 16S | 477 | 4.8 |
| 16S | 663 | 8.0 |
| 16S | 952 | 9.6 |
| 16S | 973 | 4.9 |
| 16S | 1262 | 5.9 |
| 12S | 164 | 9.0 |
| 12S | 377 | 10.0 |
| 12S | 708 | 9.0 |
| 12S | 754 | 5.0 |
| Ave |  | 7.6 |

**Figure S1**. Violin plots for the same data used to construct the 95% CI plots in Figs. 3 and 4.

**TNF-α Exposure**

**H_2_O_2_ Oxidations**

**Figure S2**. Analysis of changes in G oxidation in the 18S vs. 28S rRNAs.

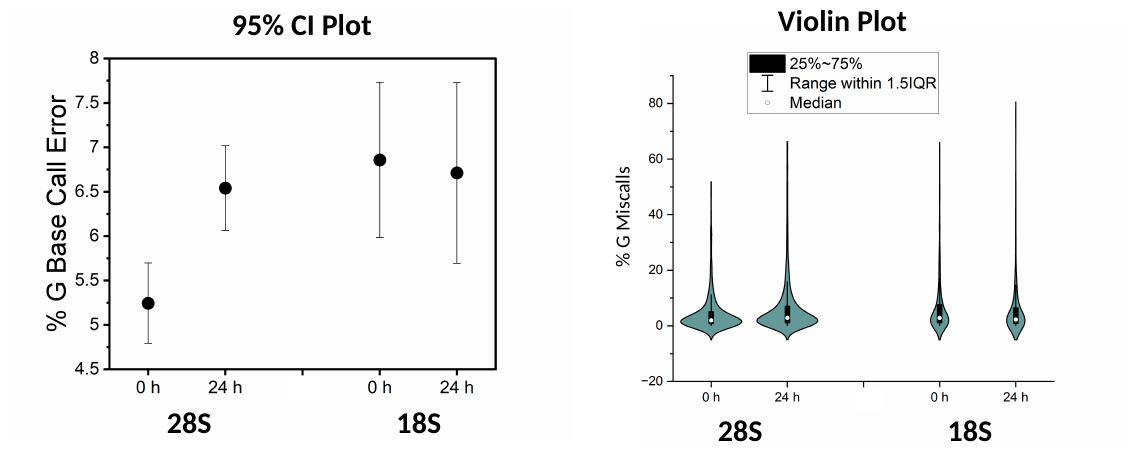

**Figure S3**. Nucleotide compositions of E. coli and human rRNAs.

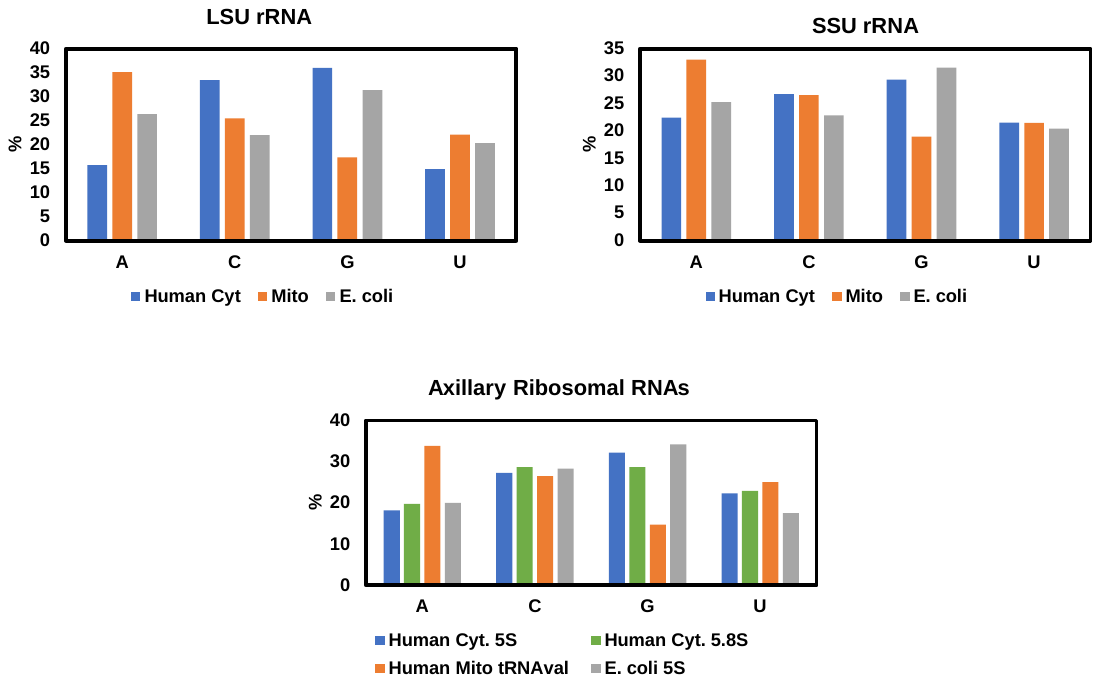

The values for the human rRNAs were obtained from the following reference sequences provided with the sequencing data. The values for the *E. coli* rRNAs were obtained from reference sequences in the literature (7).

**Figure S4**. Structure of human cytosolic ribosome with lesions mapped on the structure.

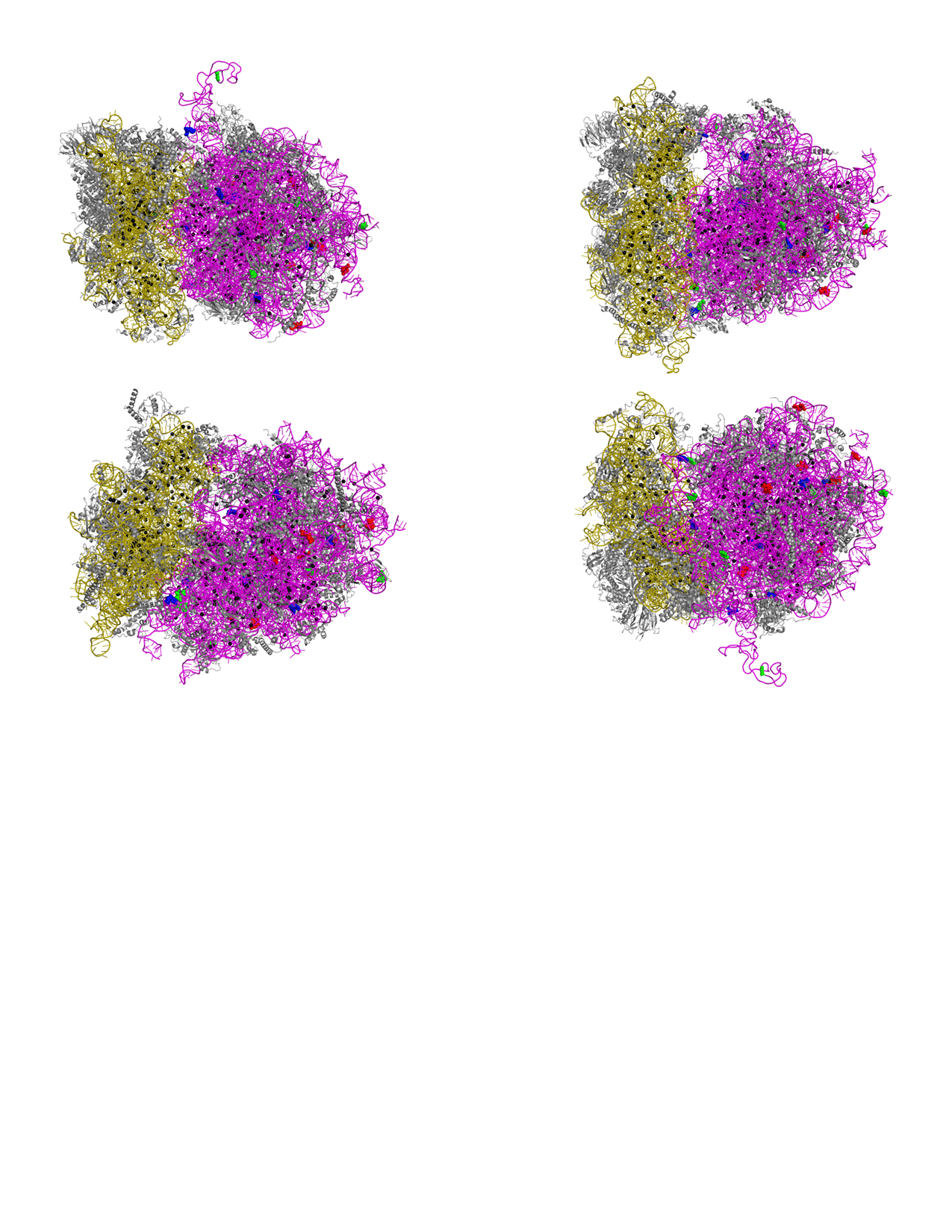

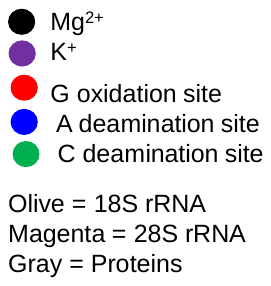

Ten G oxidation, A deamination, and C deamination sites were mapped onto the 28S rRNA to provide a visualization of where the lesion sites are on the structure. The structure pdb 6QZP was used in the construction of this figure (3).

**Figure S5**. 2D structures for sites of A-to-I deamination in the cytosolic ribosome.

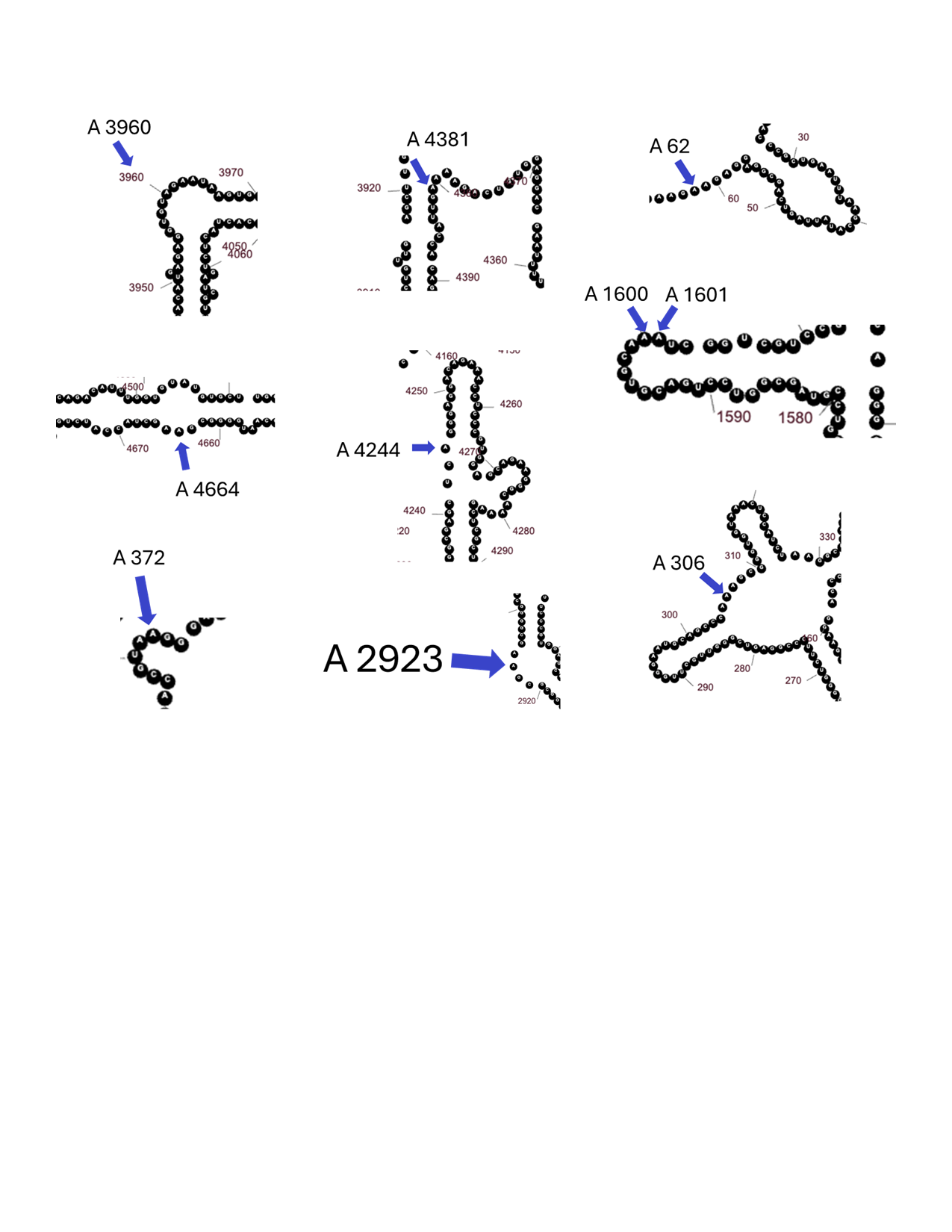

Ten A-to-I deamination sites found in the 28S rRNA are mapped onto the 2D structure of the long RNA. These sites illustrate the reactive A nucleotides are not in duplex RNA regions that are ADAR substrates, but rather in hairpin loops, bulges, or single-stranded regions where they are more solvent exposed to react with diffusible nitrosating agents.

**Figure S6**. 2D structures for sites of C-to-U deamination in the cytosolic ribosome.

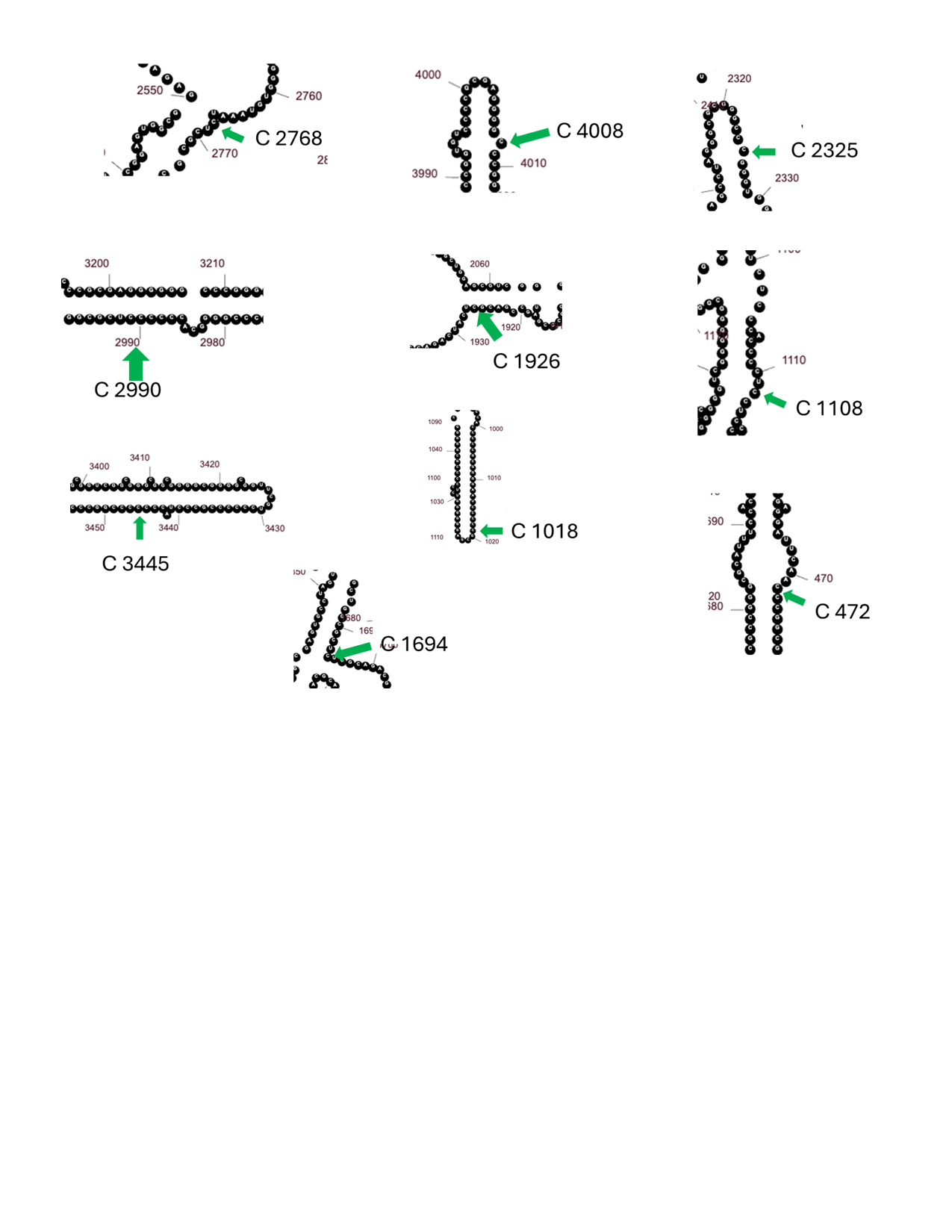

Ten C-to-U deamination sites found in the 28S rRNA are mapped onto the 2D structure of the long RNA. These sites illustrate the reactive C nucleotides are in hairpin loops, bulges, or the base of stems where they are more solvent exposed to react with diffusible nitrosating agents.

**Figure S7**. Comparative plots for the 5S, 5.8S, and 18S rRNAs before and after 24 h TNF-α exposure.

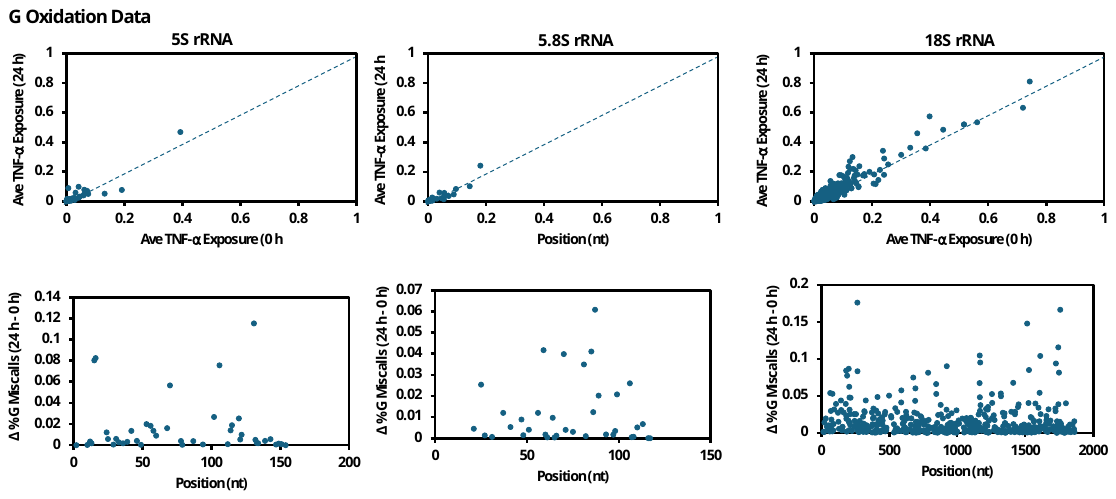

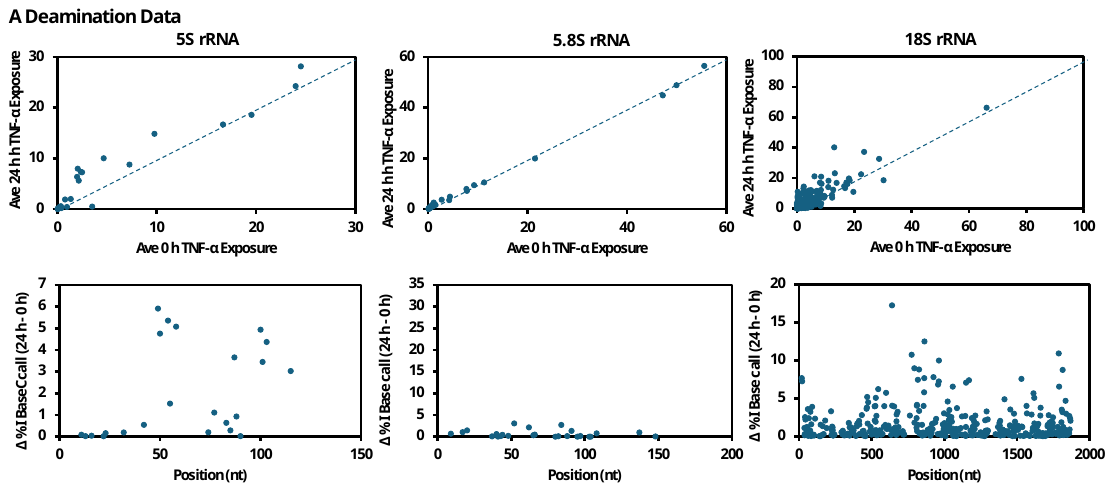

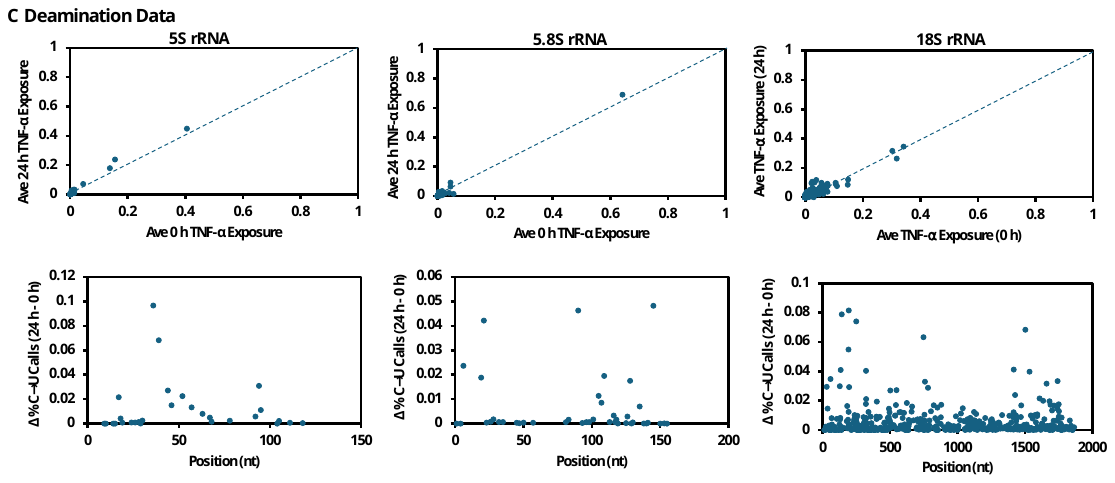

**Figure S8.** Mitochondrial ribosome with top lesion sites mapped on the structure.

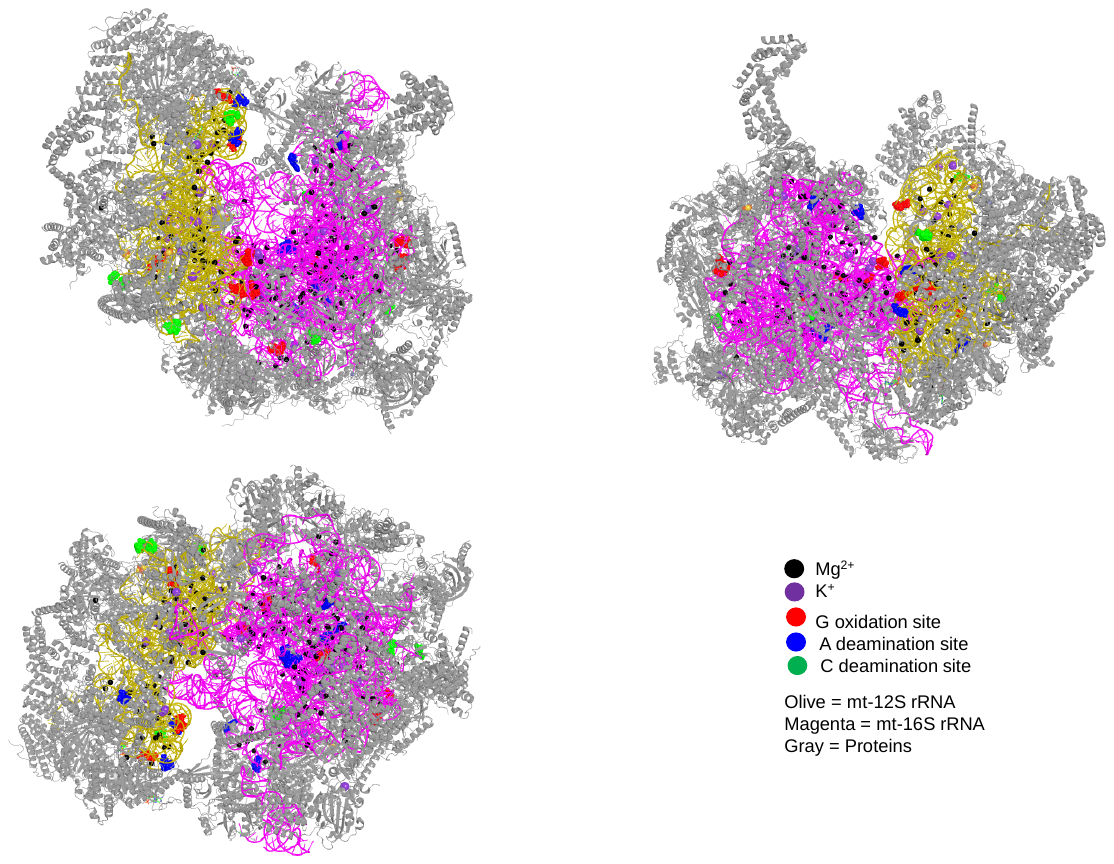

These images are provided to illustrate where the top 10 reactive nucleotides reside in the 3D structure of the mitochondrial ribosome. The structure pdb 7QI4 was used in the construction of this figure (8).
